# Clustering of SARS-CoV-2 membrane proteins in lipid bilayer membranes

**DOI:** 10.1101/2025.10.09.681538

**Authors:** Joseph McTiernan, Yuanzhong Zhang, Siyu Li, Thomas E. Kuhlman, Umar Mohideen, Michael E. Colvin, Roya Zandi, Ajay Gopinathan

## Abstract

The accumulation of viral structural proteins along the ER-Golgi intermediate compartment (ERGIC) membrane leads to SARS-CoV-2 self-assembly and budding, driven by the interactions between these proteins, RNA and the ERGIC membrane. The membrane protein (M) is believed to interact with other structural proteins and form clusters needed for the induction of membrane curvature that facilitates virion formation. However, the role played by direct and membrane-mediated interactions between M proteins and their interactions with other proteins in the clustering process remains unclear. Here, we utilize a combination of all-atom molecular dynamics (MD) simulations, continuum modeling and experiments to show that M-M interactions are sufficient to drive clustering in ERGIC-like lipid bilayers in the absence of other proteins or RNA. Using all-atom MD simulations we were able to estimate the membrane thinning induced by M proteins and the resulting membrane-mediated M-M interaction. Combining this with a continuum model that describes the evolution of M protein density in a planar lipid membrane, we identified the existence of a critical, direct M-M interaction energy needed for cluster assembly at a given density. By comparing the model predictions with analysis of atomic force microscopy images of M protein clusters in supported lipid bilayers, we were able to estimate the direct M-M interaction energy and found it to be significantly larger than the membrane mediated interaction energy. Our work therefore establishes that M protein interactions are sufficient to drive clustering and provides a quantitative understanding of the role played by direct and membrane-mediated interactions of M proteins in viral assembly and budding.

## Introduction

Each of the four prominent structural proteins in coronaviruses are required for the virus’ efficient function and replication [1–3]. As the most studied structural protein, the spike or S protein is responsible for binding the virion to the host cell, allowing its entry [1, 2, 4]. The membrane (M) protein is believed to keep the surrounding membrane of the virion stable and defines its shape [3, 5, 6], while the envelope (E) protein acts as an ion channel embedded within this membrane [7, 8]. Lastly, the nucleocapsid (N) protein recruits viral RNA and remains inside the virion [9–11]. A fully formed coronavirus virion takes a typical form, with S, M, and E proteins along the surface of a membrane surrounding viral RNA bound to N protein [1, 2]. Outside of the S protein, the other three structural proteins do not play a vital role in binding to the host cell, instead being required for efficient and complete virion formation.

Depending on the strain of coronavirus, its replication through assembly and budding along the endoplasmic reticulum golgi intermediate compartment (ERGIC) requires M protein and either N protein bound to vRNA, E protein, or both [12–15]. However, in every case, M proteins and their interactions with each other [16, 17] are required for complete virion formation [1, 3]. Furthermore, the M protein is the most prominent in every strain, and is thought to be responsible for guiding the S [3, 18], N/RNA [19–21], and E protein [3, 22] throughout the assembly and budding process. After accumulation in the ERGIC, M proteins interact with S and E proteins along the membrane surface, possibly leading to the induction of membrane curvature through protein clusters. Within the surrounding cytoplasm, N protein recruits viral RNA, which in turn binds to these M protein clusters along the surface of the membrane, believed to introduce additional curvature. After sufficient curvature generation, the M/E/S populated ERGIC membrane curls around the virus’ genetic material, leading to the budding of the ∼ 100 nm virion into the ERGIC [23–25]. Here, we focus on the initial formation of SARS-CoV-2 M protein clusters vital for viral assembly.

Recent cryo-EM experiments have found the SARS-CoV-2 M protein to be a homodimer with two different conformations, a ‘short’ compact form and an elongated or ‘long’ form [19, 26]. Cryo-ET and cryo-EM observations of the full SARS-CoV virion showed the short form in thinner regions of membrane with lower curvature as opposed to the long form [6]. Furthermore, as a result of an improved method for synthesizing the M protein, we were previously able to confirm the thinning behavior of the short form combining all-atom molecular dynamics simulations (MD) with atomic force microscopy (AFM) images [27]. We also found that M protein clusters could form within an ERGIC-like membrane depending on the protein density. However, only the short form was observed throughout the study, in agreement with previous assumptions for SARS-CoV and SARS-CoV-2 that the long form requires additional structural protein interactions [6, 26, 27] for stability.

Here, we seek a quantitative understanding of how interactions between M proteins and their density govern the formation and size of self-assembled clusters. An improved understanding of the formation of these M protein clusters is vital to clarify the expected assembly and budding of the virus. For example, it is unclear whether the thinning behavior of the M protein helps the virion clip off from the membrane, or if it plays an enhanced role in forming aggregates. The relative magnitudes of membrane mediated and direct M-M interactions and their contribution to clustering are also not known. Understanding both the effects of a single protein on the membrane and the clustering of hundreds to thousands of proteins requires a multiscale approach. In this paper, we therefore utilize a combination of all-atom molecular dynamics (MD) simulations, continuum modeling and experiments to address these unknowns.

Initially, utilizing an extended 2 *µs* all-atom MD simulation, we measured the thinning profile of the membrane in the vicinity of the M protein and computed a corresponding line tension of 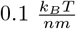 . The profile we obtain is in agreement with our earlier, shorter 1 *µs* simulations [27] and observations of the SARS-CoV M protein [6].

Next, we characterized M protein assembly within a membrane through atomic force microscopy. Following the methods described in [27] to generate AFM images of M protein embedded in a 2.25 *µm* × 2.25 *µm* suspended lipid bilayer physiologically similar to the ERGIC, we identified the existence of a critical protein density above which we saw cluster formation. We also analyzed the distribution of inter-cluster distances.

To understand the contributions of membrane-thinning induced interactions and direct M-M interactions to clustering, we turned to analytical modeling. We utilized a Cahn-Hilliard model [28, 29] on a flat plane, an approach commonly used to describe phase separation of proteins in membranes [30]. Using our model, we were able to analytically and numerically determine how these clusters form and evolve as a function of M protein area coverage and effective interaction energy. The existence of a critical effective interaction energy for each density was identified. Here, effective interaction energy encompasses all nearest-neighbor interactions experienced by an individual protein. Through direct comparison with AFM scans and our MD thinning profile determined line tension, we estimated the effective interaction energy of the M protein as *ϵ*_*m*_ ∈ [7.8*k*_*B*_*T*, 9.6*k*_*B*_*T* ] and the direct M-M interaction as *ϵ*_*direct*_ ∈ [6.7*k*_*B*_*T*, 8.5*k*_*B*_*T* ]. As a result, we inferred that membrane thinning does not play a substantial role in M protein cluster formation and is more likely to be prominent elsewhere in the assembly and budding process. Additionally, the density fraction needed for these assembly-like formations in a flat plane is [0.118, 0.304]. This average density range and the existence of a critical effective interaction energy given an average density suggests possible methods for inhibiting cluster formation. Inhibition of cluster formation prevents the first step of the assembly and budding process, helping suppress SARS-CoV-2 replication.

## Materials and methods

### All-Atom Molecular Dynamics

The all-atom molecular dynamics (MD) simulation of the short form embedded in a lipid bilayer was performed using the CHARMM36m force field with the MD package GROMACS, version 2022.3 [31, 32]. The CHARMM-GUI input generator was used to set up the simulated system with periodic boundary conditions, and supplied the six steps used for equilibration [33–41]. After equilibration, the system was simulated for 2 microseconds with a timestep of 2 femtoseconds in the NPT ensemble. System temperature was maintained at 303.15 K using the Nose-Hoover thermostat [42, 43], with the pressure maintained semi-isotropically at 1 bar in the x-y dimensions and separately in the z-dimension using the Parrinello-Rahman barostat [44, 45]. The coordinates were saved once every 50 thousand timesteps, or every 0.1 ns, for a total of 20 thousand frames.

The protein was inserted into the membrane with an orientation and depth that matched other studies [19, 26]. This insertion can be seen in Fig. 2a and 2b. Only residues 9-204 are accounted for in the short form structure (PDB: 7vgs), with the first eight and last 18 residues excluded [19]. The membrane was composed of Chol 15%; DOPC 45%; DOPE 20%; DOPS 7%; POPI 13% in both leaflets, with the solvent consisting of NaCl at a concentration of 0.15 M and TIP3P water. Initially, the simulation box consisted of a 24.7 nm x 24.7 nm membrane in the x-y plane with at least 5 nm of solvent above and below the protruding protein, yielding a total unit cell thickness of 17.1 nm in the z dimension. However, by the end of the simulation, the membrane size changed to 24 nm x 24 nm, with a z-axis box size of 18.1 nm. Atom counts and box size evolution can be seen in S1 Fig. The final trajectory was reoriented frame-by-frame such that the protein was centered in the box for all frames. While each trajectory was fitted to eliminate protein translation, this was not the case for the rotation of the protein. All-atom simulations were visualized using ChimeraX [46], with the python library MDAnalysis used to analyze the processed trajectory [47, 48].

Membrane thickness, defined in Fig. 1c, is computationally determined by creating an 18 x 18 square grid in the x-y direction with edges defined by the minimum and maximum phospholipid head in either direction. At each point in time, these heads are binned, and the average height of the lower and upper leaflets are determined. In the case a bin does not have any head atoms, the corresponding upper and lower leaflets are ignored for that position at the given time. From here, the difference between average height for each leaflet is calculated at every point in time beyond 500 ns, and averaged over time to get the figure shown in Fig. 1d.

**Fig 1.**
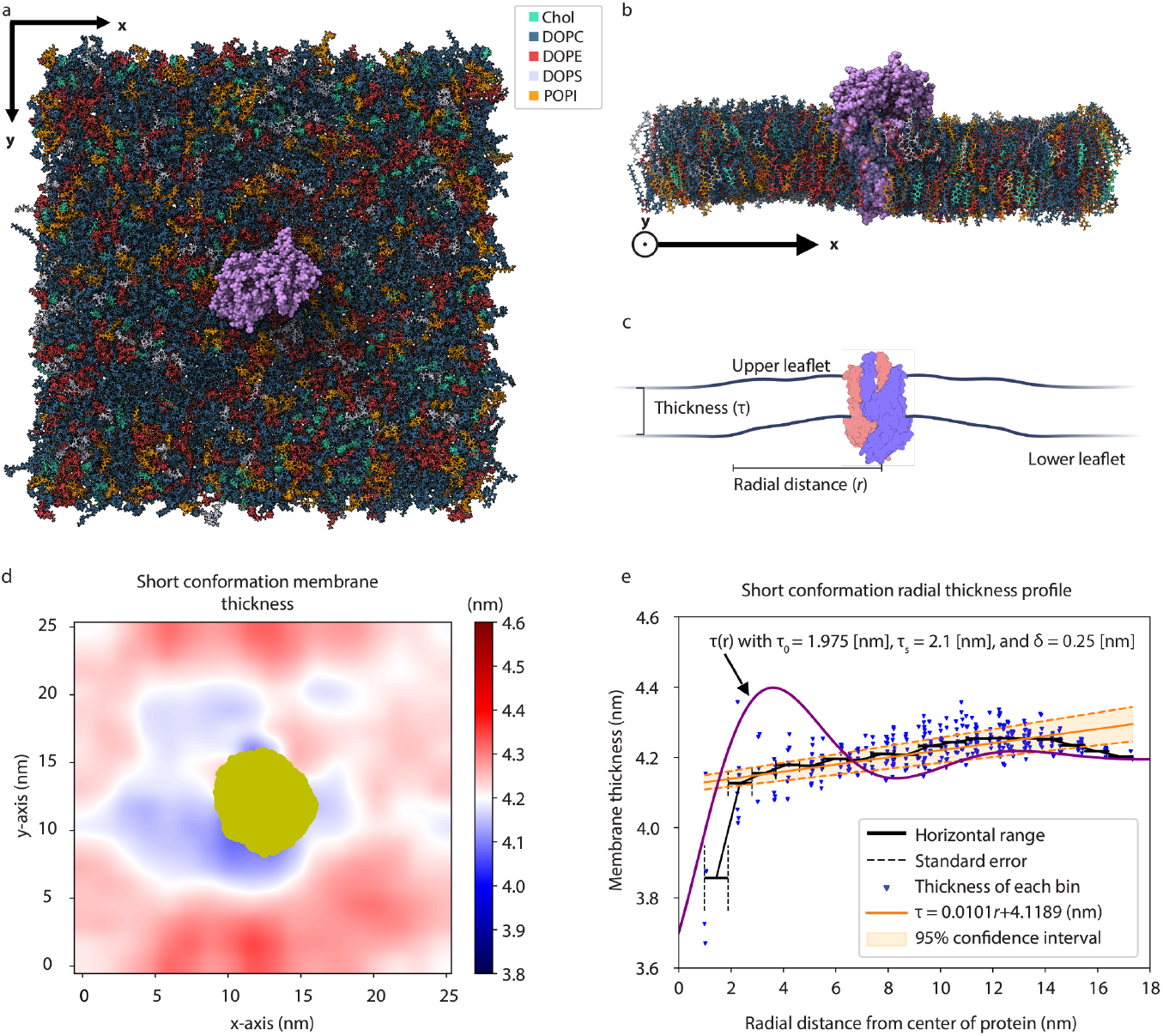
All-atom molecular dynamics of the M protein short form embedded in a multicomponent membrane. Final simulation frames of the short form M protein embedded (purple) in a 25 [nm] x 25 [nm] lipid bilayer physiologically similar to the ERGIC from above (a) and as an x-axis cross-section (b). (c) Cartoon defining membrane thickness (*τ* (*r*)) as the difference between the upper and lower leaflets for a given radial distance *r*. The two chains in the M protein are distinguished with red or blue, where the C-terminal of the protein pointing downwards is within the virion. (d) Average membrane thickness is shown along the x-y plane from 500 [ns] to 2000 [ns], where red represents thicker regions of membrane and blue thinner. Gold signifies the cumulative cross-section of the protein. (e) Radial thickness profile relative to center of protein, obtained from averaging (d) over all angles, is shown with blue representing individual bins. A 95% confidence interval for a line of best fit is shown in orange, with the black trend line representing a radial binning of the blue points. The purple curve represents the analytic solution to the thickness profile given with *τ*_0_ = 1.975 [nm], *τ*_*s*_ = 2.1 [nm], and *δ* = 0.25 [nm].

**Fig 2.**
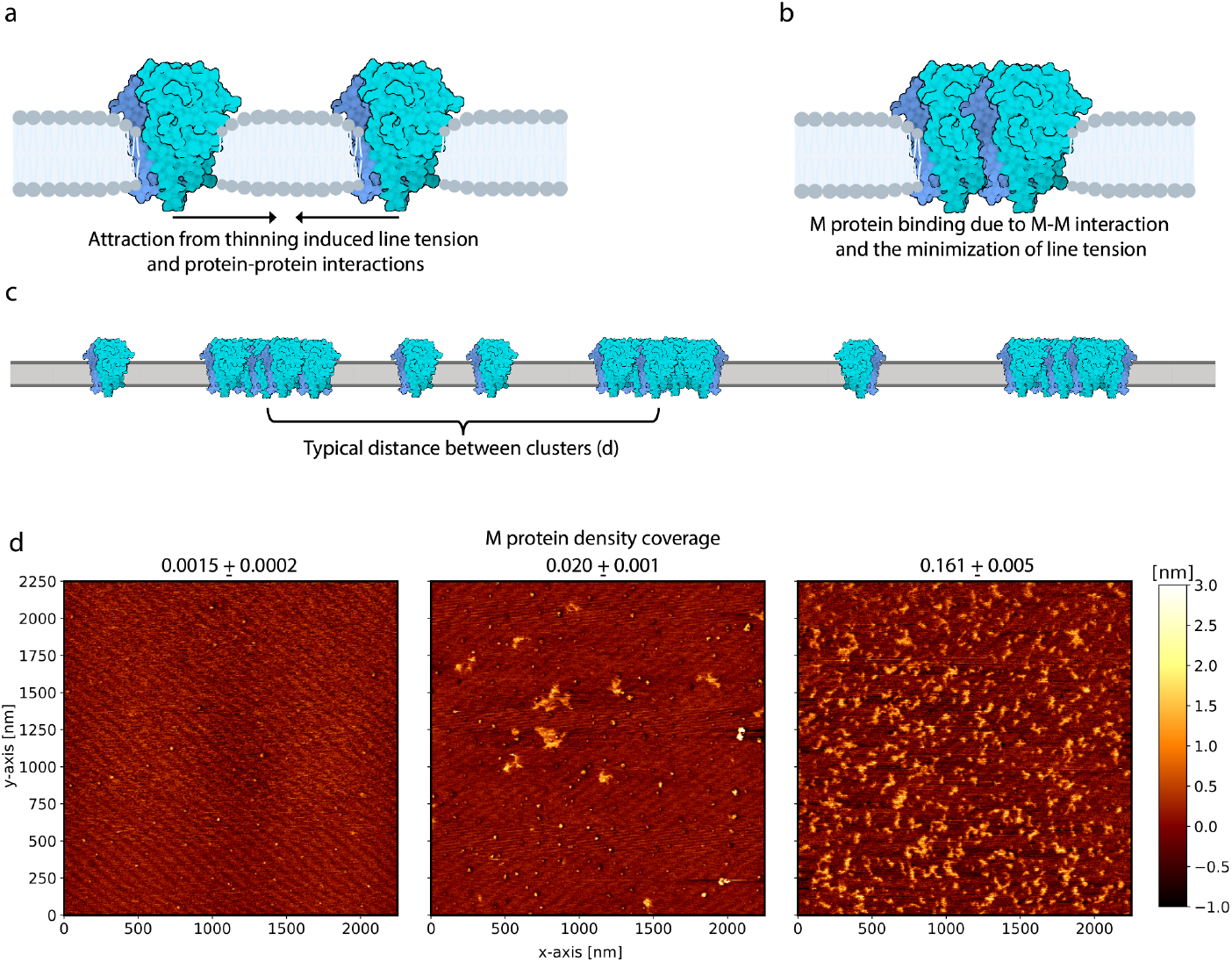
Atomic force microscopy height images of 2.25 *µm* × 2.25 *µm* suspended lipid bilayer showing cluster formation at different M protein area coverages. (a) Cartoon showing nearby M protein attracting as a result of minimizing membrane thinning induced line tension. Larger range protein-protein interactions could also help this process. The different shades of blue represent the monomers within the dimer. (b) After attracting an adjacent protein, they will bind together as a result of direct M-M interactions and to further minimize line tension. (c) With sufficient protein density, enough of these complexes will form to create a periodic arrangement of clusters, defined by an average distance between them (*d*). (d) AFM height images of a 2.25 *µm* ×2.25 *µm* suspended membrane for three different M protein mass to lipid mass ratios. The corresponding protein density area coverage is shown for each ratio, increasing from right to left, where lighter regions represent greater heights.

### Thinning Induced Line Tension

Assuming a symmetric deformation, the membrane thickness profile can be considered the combination of two equivalent monolayers. Adapting the model developed in [49] for determining monolayer height in a transition region between higher and lower membrane thickness due to lipid rafts leads to Eq. 1. In this case, the protein is considered a raft with very high elastic moduli which deforms the surrounding membrane to match its hydrophobic thickness. This expression provides membrane thickness as a function of distance from the protein (*r*), where *τ*_*s*_ is the thickness of the unperturbed monolayer, *τ*_0_ is the average between low (*τ*_*r*_) and high thickness monolayers, *δ* is the difference between these regions (*δ* = *τ*_*r*_ − *τ*_*s*_), 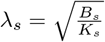 and 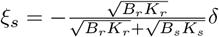. The factor of two in the equation converts thickness to that of a bilayer, and the contribution from induced curvature is taken to be negligible.

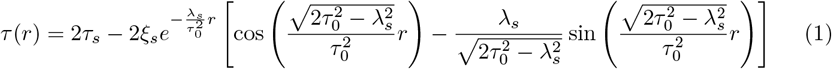

In [27] we showed, using AFM methods, that regions with protein are much stiffer than the surrounding membrane. With *B*_*s,r*_ and *K*_*s,r*_ the bending and tilt moduli of the surrounding membrane and the raft/protein respectively, *B*_*r*_ *>> B*_*s*_ and *K*_*r*_ *>> K*_*s*_. Furthermore, [27] gives *B*_*s*_ ∼ 3 *k*_*B*_ *T* and 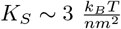, leading to *λ*_*s*_ ∼ 1 nm and *ξ*_*s*_ ∼ −*δ*.

With this thickness profile, the line tension generated from the cost of bending and tilt can be determined as described in [49]. Without the negligible contributions from induced curvature, Eq. 2 shows the line tension *γ*_*m*_. Where *B*_*r*_ *>> B*_*s*_ and *K*_*r*_ *>> K*_*s*_ allows for the corresponding approximation, and the factor of two accounts for the bilayer nature of the membrane.

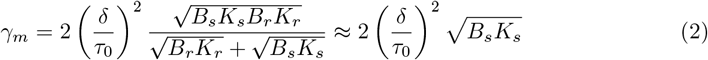

### Protein Assembly Continuum Model

To properly represent the assembly of a large number of M proteins along a flat membrane, we utilize a continuum model. We use a Cahn-Hilliard model [28, 29] to describe the protein density evolution on a flat plane, an approach commonly used to describe phase separation of proteins in membranes [30]. It is to be noted that such approaches have also been extended to describe the motion of curvature inducing transmembrane proteins [30, 50–52], though here we consider only the planar lipid bilayer case and do not consider density dependent membrane surface tension nor membrane viscosity [51–53]. The total free energy of the system consists of an enthalpic contribution arising from protein-protein and protein-membrane interactions and an entropic part,

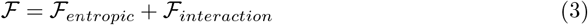

The entropic portion of the free energy is a typical entropy of mixing which can be expressed as,

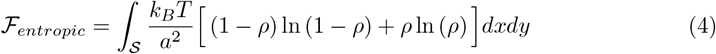

where *ρ* is the protein density fraction which is a two dimensional scalar field along a flat infinitesimally thin surface, *T* is the temperature, and *a* represents the nearest neighbor distance between proteins.

The interaction portion of the free energy, ℱ _*interaction*_, involves three different terms,

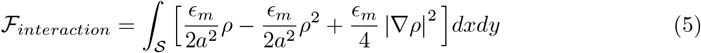

where *ϵ*_*m*_ is an effective interaction energy accounting for all nearest neighbor interactions a single M protein would experience (refer to Effective Interaction Energy for details) [54]. An effective interaction energy greater than zero implies attraction. This has the form of a two-component regular solution or Flory-Huggins model [28, 54–56], with volume exclusion, protein attraction, and interfacial energy terms.

Following a standard approach [30, 51], the dynamics of *ρ* can now be expressed using Model B dynamics to account for the conservation of protein density leading to

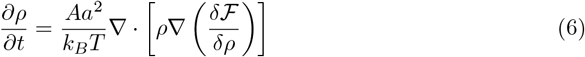

The full expression for the variational derivative 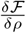 (Eq. 18) and full density evolution equation (Eq. 19) can be seen in S1 Appendix.

### Linear Stability Analysis

To quantitatively understand the conditions and parameter regimes required for clustering, we performa linear stability analysis. This is done by first applying a small perturbation to a homogeneous density state of the form

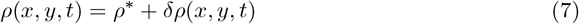

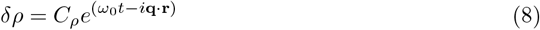

Here, the initial protein density fraction is *ρ*^∗^, and the perturbation *δρ <<* 1 (set by *C*_*ρ*_ *<<* 1). Each mode is defined by its wavevector **q** and growth rate *ω*_0_. Throughout this work, **q** is nondimensionalized according to 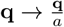. A positive growth rate for any *q* implies an unstable mode with that wavevector and indicates clustering with an average distance between cluster formations (*d*) set by 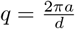 . A schematic characterizing protein aggregation in an unstable regime can be seen in Fig. 3.

**Fig 3.**
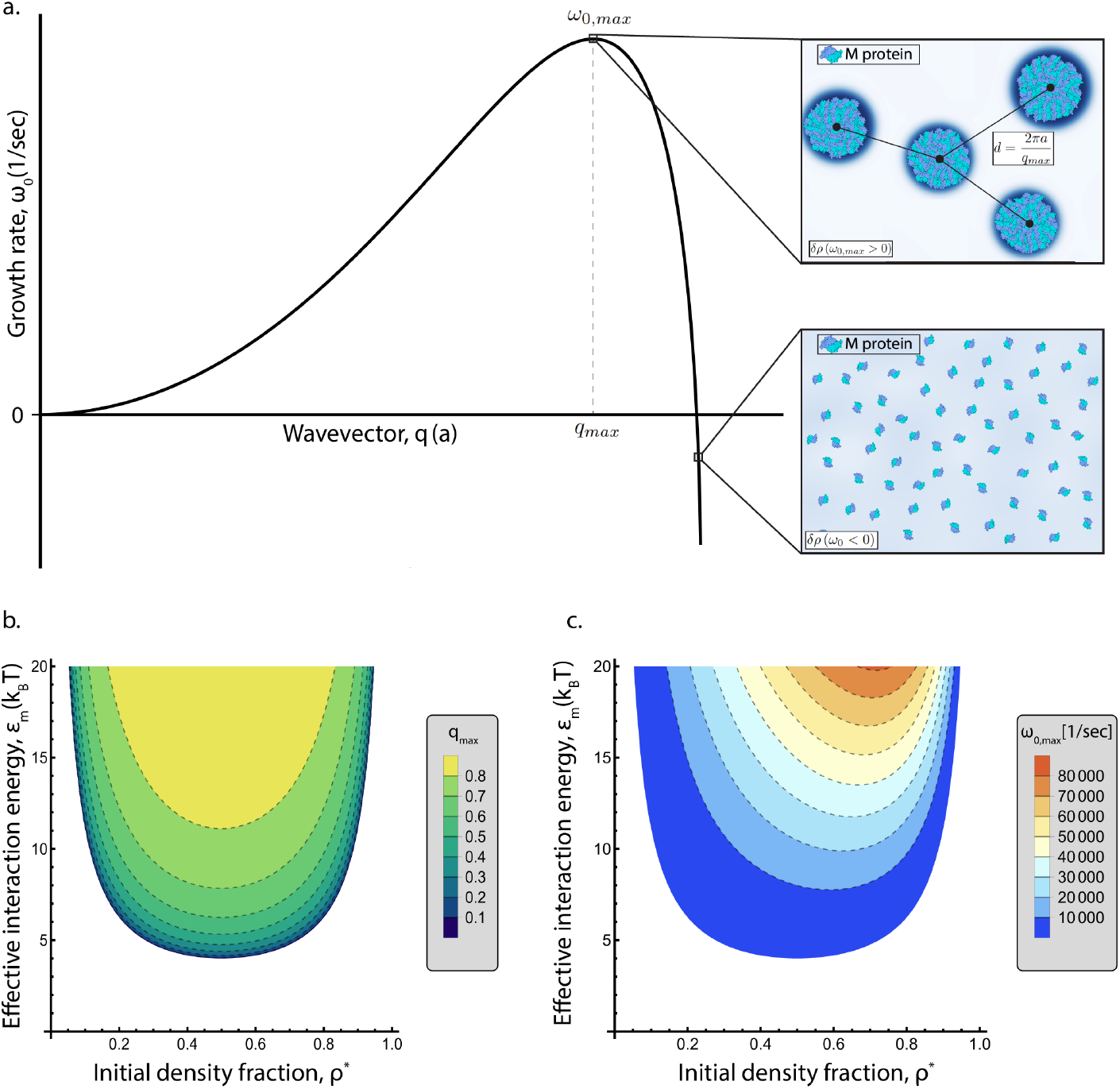
Linear stability analysis of M protein aggregation. (a) Typical unstable dispersion relation, where each mode for the perturbation *δρ* is defined by its wavevector *q* and its growth rate *ω*_0_. A growth rate greater than zero represents an unstable mode, where the fastest growing mode is described by *ω*_0,*max*_. Within this mode, M protein clustering will occur for an average distance between clusters *d*, as seen in the upper right panel. A stable mode is shown in the bottom right panel, where proteins remain isolated. Phase diagrams for maximum wavevector (b) and maximum growth rate (c) with respect to the effective interaction energy (*ϵ*_*m*_) and the initial protein density fraction (*ρ*^∗^). Each dotted grey line represents a corresponding contour in the provided legend.

Using Eq. 7 in Eq. 6 and ignoring all nonlinear terms leads to the protein evolution equation

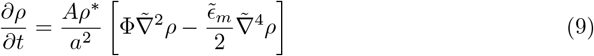

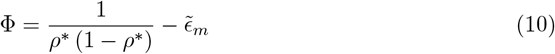

where tildes signify nondimensionalized quantities 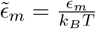 and 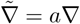 ). Using the form of the perturbation in Eq. 8 in Eq. 9 allows us to obtain the growth rate as a function of wavevector

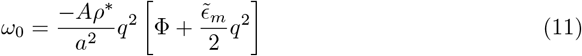

From this dispersion relation, the wavevector and growth rate corresponding to the fastest growing modes can be determined.

### Effective Interaction Energy

The effective interaction energy (*ϵ*_*m*_) (or Flory Huggins interaction parameter) is determined by considering a lattice where lattice sites can be occupied by membrane or a protein and only nearest neighbor and spatially independent interactions are considered. *ϵ*_*m*_ can then be expressed in terms of membrane and protein interactions as [54]

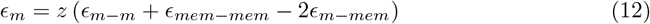

where *z* defines the number of nearest neighbors each site has, and *ϵ*_*i*−*j*_ is the interaction between two sites occupied by entities of type *i* and *j* (membrane (mem) or protein (m)) a distance *a* apart where *i, j* ∈ {*m, mem* }.

Each site-site interaction term can be determined using smaller scale MD simulations [57]. However, in doing so, numerous unknown parameters and membrane characteristics are introduced to the system, ranging from lipid tilt to protein induced curvature. Furthermore, the scale and nature of AFM data prevents the analysis of these quantities. We therefore treat *ϵ*_*m*_ as an effective parameter to be estimated.

We note that the energy difference between a discrete lattice with two proteins spread apart and one with them adjacent to each other is equivalent to Eq. 12. This energy difference can also be considered the energetic cost resulting from line tension and direct protein-protein interactions. As a result, the effective interaction energy can be expressed as,

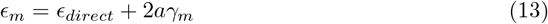

where *γ*_*m*_ is the thinning induced line tension and *ϵ*_*direct*_ is the energy in which these proteins are bound together. A more detailed derivation of this concept is shown in S2 Appendix.

### Finite Difference Method

To numerically solve the protein evolution equation shown in Eq. 6 and compare with the analytical cluster formation predictions, a finite difference method is utilized. The python code developed for this finite difference method can be found at https://github.com/jmctiernan24/M_Protein_Assembly. Spatial derivatives are determined through fourth order central difference, while Fourth Order Runge Kutta is used for time evolution. Membrane (M) protein is initially distributed normally with an average equal to the initial protein density fraction (*ρ*^∗^) and a standard deviation of 5 × 10^−5^, over a 250 x 250 grid. Each grid point is 0.4*a* in length, where *a* is the size of an M protein and is assumed to be *a* = 5 nm, while corresponding time steps depend on the effective interaction energy within the range of [1 × 10^−9^, 5 × 10^−9^] [sec]. High spatial and temporal resolution is required due to the lack of a more sophisticated finite element method. The temperature used throughout every simulation is *T* = 303.15 *K*, with the diffusion constant 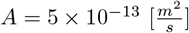 .

To compare with the analytically determined wavevector and growth rate, a radial power spectrum of the density is calculated for every 100 time steps within linearity, defined as *δρ <* 0.01. The maximum wavevector (*q*_*max*_) is determined by fitting a Gaussian to the radial power spectrum, and equating it to the point in which there is a maximum. From here, the square root of this Gaussian at the maxima for each measured time step is fit to exponential growth.

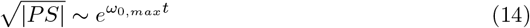

Example power spectra and fits are shown in S3 Fig. Outside of linearity, the nearest neighbor distance is used to classify cluster formation periodicity. Clusters are defined according to a density threshold such that the protein area fraction is equivalent to the initial protein density (*ρ*^∗^). This process can be seen in S4 Fig for each of the images shown in Fig. 4b. The evolution of nearest neighbor distance can be seen in S5 Fig, and is measured every 10,000 time steps upon *ρ*_*max*_ *>* 0.9. Time cut-offs for calculating nearest neighbor distance to minimize the impact of Ostwald ripening were found by determining the flattest region of the nearest neighbor distance evolution plots, with the cut-offs shown in S5 Fig.

**Fig 4.**
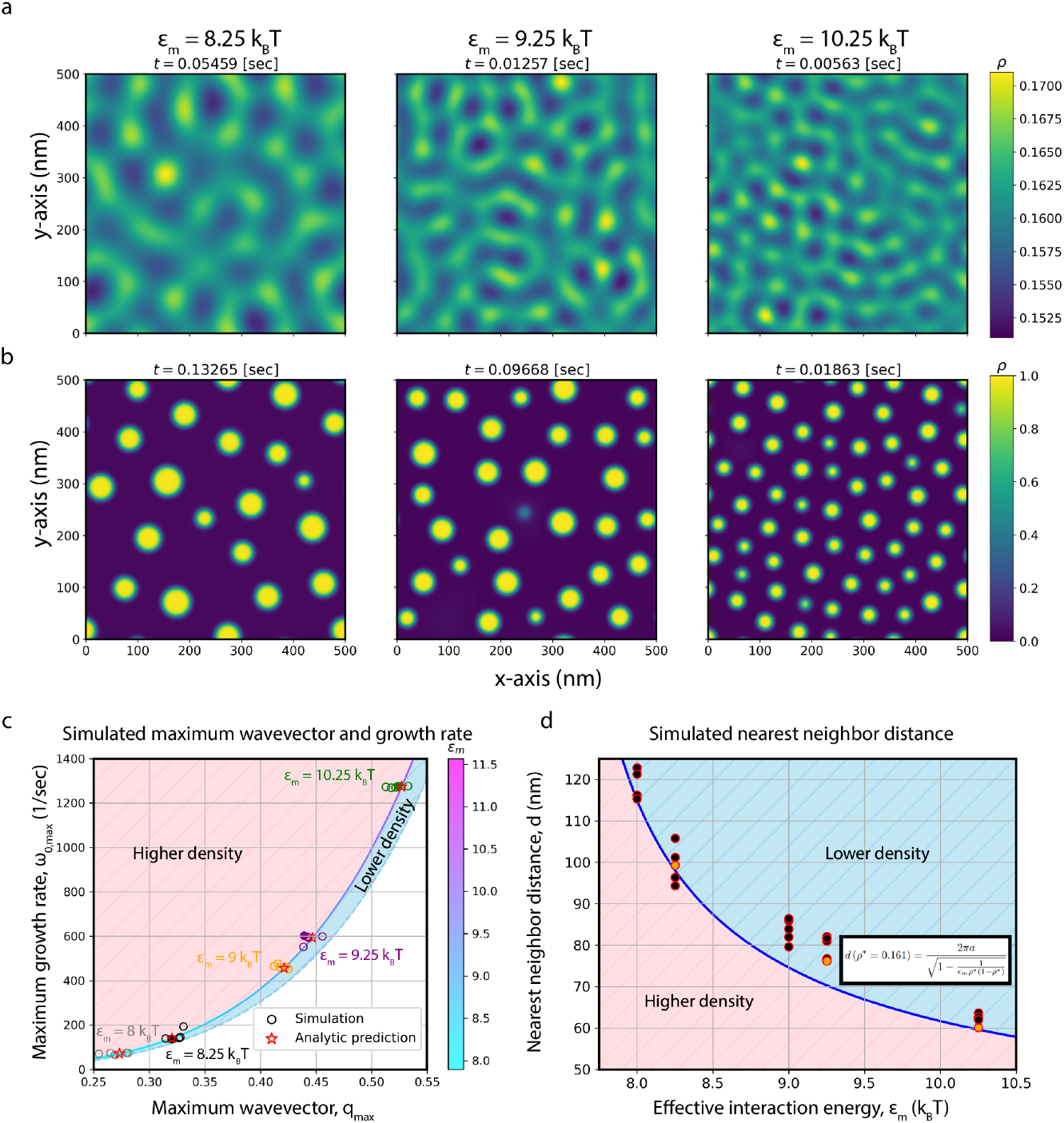
Evolution of M protein density and clustering within and beyond linearity. Simulations for three different interaction energies are shown for *ρ*^∗^ = 0.161, with (a) showing each at the final frame of linearity and (b) displaying later frames of a stable state before Ostwald ripening begins to dominate. The time of the frame is shown above the image, with brighter regions representing higher density, where the scale of the colorbar changes between (a) and (b). (c) Plot of the numerically determined maximum growth rate and maximum wavevector for the shown simulations and additional replicates within linearity. Each interaction energy is identified with a different color, where replicates are shown as hollow circles, those shown in (a) and (b) are filled in circles, and the analytic prediction for a given interaction energy is a red star. The analytic prediction for each interaction energy is shown with the line between the higher/lower density regions, which changes color according to *ϵ*_*m*_ in the corresponding colorbar. (d) Comparison between average nearest neighbor distance at stable states of numerical simulations to an analytic prediction for different interaction energies. Simulations shown in (a) and (b) are filled in orange, while extra replicates are filled with black. Representing the analytic prediction, the blue curve is determined using the highlighted equation. For (c) and (d), the striped red region represents the direction the respective prediction would move if increasing density, while the striped blue region gives the reverse.

### AFM Sample Preparation and Imaging

The methods in this section are similar to those from [27]. First, monodisperse LUVs around 120 nm in diameter were prepared using a vesicle extruder, with lipid composition corresponding to that of the endoplasmic reticulum-Golgi intermediate compartment (ERGIC). The molar ratio of different lipids (Avanti Polar Lipids, Inc, Alabaster, AL) used are 1-palmitoyl-2-oleoyl-glycero-3-phosphocholine (POPC): 1-palmitoyl-2-oleoyl-sn-glycero-3-phosphoethanolamine (POPE): 1-palmitoyl-2-oleoyl-sn-glycero-3-phosphoinositol (POPI):1-palmitoyl-2-oleoyl-sn-glycero-3-phospho-L-serine (POPS): Cholesterol = 0.45: 0.2 : 0.13: 0.07: 0.15 [58]. The 5 mg/ml lipid solution in chloroform was dried in a glass vial with N2 gas stream, then vacuumed overnight at -30 in Hg. The dried lipid mixture was hydrated with buffer (150 mM NaCl, 20 mM HEPES, pH = 7.2) and 30 s vortex, prior to ten freeze-thaw cycles with dry ice and 37 °C bath. After the final thawing step, the aqueous solution was passed 11 times through a polycarbonate membrane with 100 nm pores (Nuclepore Track-Etch membrane, Whatman, Chicago, IL).

To reconstitute the M proteins into LUVs, concentrated stock solution ( ∼ 400 mM) of n-dodecyl-*β*-D-maltoside (DDM Avanti Polar Lipids, Inc, Alabaster, AL) were added into 5 mg/mL of freshly extruded LUVs solution to reach a final concentration of 100 mM. M-protein stabilized by Triton X-100 (stock solution: 1 wt.% Triton X-100 per every 2 mg/mL M protein) was added to the LUVs solution at mass ratio of M/lipid = 1/100, 1/67, and 1/50 after 10 min of incubation. The solution was allowed another 10 min of incubation and the detergent was removed using 80 mg of wet BioBeads (Bio-Rad, Hercules, CA) per mL of LUVs. After 3 additions of BioBeads (once per 2 hours), the M-reconstituted LUVs were separated from the solution using the centrifuge.

For the preparation of supported bilayer samples for AFM imaging, 75 *µ*L of the solution containing LUVs collected from the bottom of a microcentrifuge tube was deposited onto freshly cleaved pristine mica and incubated for 1 hour. Next, the sample was then gently rinsed with 5 mL of buffer. The samples were kept submerged by buffer during the sample preparation and imaging process. Imaging was performed within 1 hour of sample preparation.

An AFM fluid cell filled with buffer with an MSNL cantilever (Bruker, Camarillo, CA) in tapping mode was used. The spring constant was calibrated to be 0.30 N/m using the thermal oscillation spectrum. 512 x 512 pixel images 2.25 *µ*m x 2.25 *µ*m were taken, with a vertical resolution of 0.01 nm and horizontal resolution of 4.4 nm. The cantilever tip size and image resolution was calibrated using 10 nm gold spheres (Ted Pella, Redding, CA) using our previously reported [27, 59].

### Atomic Force Microscopy Cluster Determination

To define regions with and without protein from the AFM height images shown in Fig. 2, a total variation denoising filter was used (TV Bregman from the python skimage library). The impact of this filter can be seen in S6 Fig. The TV Bregman filter allows for the maintainment of sharp height changes as the probe transitions from membrane to protein, while also reducing noise within protein clusters or large regions without protein [60, 61]. After filtering the data for the case of Fig. 2, regions with or without protein are decided based off a thresholding value defined using Otsu’s method [62].

Anything above this value is a pixel with protein, while values below represent membrane. However, for the lowest area fraction, Otsu’s method is not sufficient for protein determination, and thus the threshold value for the middle density fraction is used. Refer to S7 Fig for more information.

Upon thresholding the filtered data, a cluster is defined as more than four adjacent pixels with protein. This restriction represents anything more than a single protein, since each pixel is approximately 4.4 nm across. After this restriction, the center of ‘height’ for each cluster is determined. This additional weighting accounts for proteins being off-center, or the pixel only showing the edge of the protein. Lastly, nearest neighbor distance is calculated such that each centroid is supplied a minimum distance representing the closest cluster using the sklearn python library NearestNeighbor function. For two centroids that are both closest to each other, only a single value is accounted for. To determine whether the centroid density is roughly constant throughout the images, kernel density estimation from the sklearn python library is used (KernelDensity function). See S8 Fig for centroid density images at each protein area fraction.

## Results

### M Protein Induced Thinning Profile and Line Tension

To determine the short form’s impact on membrane thickness we perform all-atom MD simulations of the M protein in its short form configuration embedded within a 25 nm x 25 nm lipid bilayer physiologically similar to the ERGIC. Final simulation frames, after 2 *µs*, can be seen in Fig. 1a and Fig. 1b, where the thickness is defined as the difference between the height of the upper and lower leaflets (Fig. 1c). Averaged over the last 1.5 *µs* of the simulation, the membrane thickness is shown from above in Fig. 1d, with the cumulative cross-section of the protein in gold. While not completely axisymmetric, the membrane is noticeably thinner near the protein. Furthermore, taking a radial profile of membrane thickness by averaging over every angle with respect to the center of the protein leads to Fig. 1e. Every bin is represented as a blue triangle, with the black trend line corresponding to averaging bins over a radial range. Additionally, the linear line of best fit and the corresponding 95% confidence interval in orange shows the significance of the short form’s membrane thinning. Similar to the shorter MD results and AFM profile displayed in [27], the short form thins the membrane by 0.5 nm starting 12 nm from the center of the protein. Confirmation of the protein’s stability can be seen in S9 Fig.

We can fit the membrane thickness data to Eq. 1, which represents an analytic prediction of the membrane thickness as a function of distance from the center of the protein (purple line in Fig. 1e). In our case, the thickness profile is parametrized by average monolayer thickness *τ*_0_ = 1.975 nm, unperturbed monolayer thickness *τ*_*s*_ = 2.1 nm, and total thinning *δ* = 0.25 nm. With the symmetric behavior of the membrane, each of these quantities are half of the full bilayer values. Integrating the elastic energy cost of this membrane thinning profile provides an estimate of the line tension, as described in Thinning Induced Line Tension. Using appropriate elastic constants for the membrane 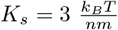 and 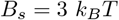 ), the line tension from bending and tilt is approximately 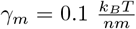 . We note that this line tension could serve as a possible membrane mediated interaction to create clusters.

### M Protein Assembly Depends on Initial Density Fraction

Next, we use AFM to directly observe M protein cluster formation. First, shown in Fig. 2a’s cartoon, two nearby M proteins can attract solely in an attempt to minimize the elastic deformation of the membrane from thinning. As these proteins attract each other, Fig. 2b displays a cartoon of the minimized line tension and role of direct M-M interactions in binding them together. Through these membrane mediated interactions, protein clusters can form (depending on the protein density) with some characteristic distance between them (cartoon shown in Fig. 2c).

Using three different protein-lipid mass ratios provides three different protein area coverage values, shown in AFM images from Fig. 2d of a 2.25 *µm* × 2.25 *µm* membrane with protein spread throughout. Area coverage is considered as the percentage of pixels occupied with a protein, where protein occupation is defined based on a threshold AFM height value dependending on mass ratio. A mass ratio of 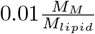 leads to the lowest area coverage, shown in the first panel of Fig. 2d, ranging from 0.0013 to 0.0017 when accounting for the 0.01 nm vertical resolution of the AFM scan. In this case, M proteins are found individually or as small oligomers without any cluster formation. As this ratio is increased to 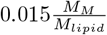 in Fig. 2d panel 2, the area coverage increases to 0.020 ± 0.001. Furthermore, larger clusters begin to form while individual proteins or small oligomers remain isolated, leading to a non-isotropic protein density. When compared to the highest mass ratio of 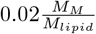, the protein area coverage jumps to 0.161 ± 0.005, shown in Fig. 2d panel 3. Within this area coverage protein clusters form roughly isotropically, where individual proteins are very rare, allowing for the characterization of a typical distance between clusters (*d*). Thus, there exists a critical M protein density between the area coverages shown in Fig. 2d panels 2 and 3, where clusters begin forming consistently with a characteristic distance. Position dependent centroid density for each of the three AFM images is shown in S8 Fig.

### Effective Interaction Energy Defines Onset of Cluster Formation

To quantitatively understand the cluster formations in our AFM experiments and to better characterize M protein assembly overall, we utilize a Cahn-Hilliard model accounting for protein interactions through two-component regular solution theory described in Protein Assembly Continuum Model. Linear stability analysis is used to identify the onset of cluster formation providing parameter regimes in which these formations are possible (refer to Linear Stability Analysis).

Briefly, we derive a dispersion relation between the wavevector *q* and corresponding growth rate *ω*_0_ of a perturbation. Unstable modes will have large positive values for *ω*_0_ and stable modes will have negative values. An example unstable relation is shown in Fig. 3a, where there exists a maximum growth rate *ω*_0,*max*_ at wavevector *q*_*max*_. This is considered the fastest growing mode and will dominate the others numerically and experimentally, leading to cluster formation with a spacing defined by the maximum wavevector, as seen in the upper right panel of Fig. 3a. Additionally, a stable mode will decay back to the initial condition, or the proteins will be spread randomly (leftmost panel of Fig. 2d and the bottom right panel of Fig. 3a).

The effective interaction energy and initial protein density fraction define the onset of cluster formation. Initial M protein density fraction *ρ*^∗^ is equivalent to the AFM area coverage, ranging from 0 - 1. Figure 3b shows these parameters’ impact on the maximum wavevector, where white represents a pairing without any cluster formation. As effective interaction energy is increased so is the maximum wavevector. Additionally, Fig. 3c shows the maximum growth rate as a function of *ρ*^∗^ and *ϵ*_*m*_, which is asymmetric about *ρ*^∗^ = 0.5 as opposed to the maximum wavevector.

Thus, for any *ϵ*_*m*_, there exists a range of *ρ*^∗^ where clusters will start to form. Furthermore, for any initial density fraction there exists a critical effective interaction energy at which the onset of assembly begins. This is in agreement with Fig. 2d, where there exists some critical density at which consistent clusters begin to form.

Inverting the curves displayed in Fig. 3b leads to a relation defining the effective interaction energy as a function of the initial protein density fraction and the maximum wavevector

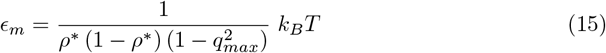

This can also be used to express the expected distance between clusters for a given interaction strength

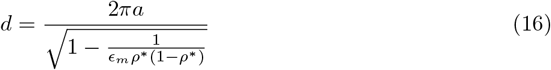

As a result, effective interaction energy can be estimated from our experimental AFM images shown in Fig. 2d, assuming the relation holds beyond linearity. See S1 Appendix for the expanded version of maximum growth rate and maximum wavevector. Solely from the ability to form clusters at *ρ*^∗^ = 0.161, a rough lower bound for the effective interaction energy can be identified: *ϵ*_*m*_ ∼ 7.4 *k*_*B*_*T* . However, direct comparison with AFM images using Eq. 15 is required for a more accurate estimation. While the onset of cluster formation is needed for further aggregation, it is unknown whether these formations will survive beyond linearity. Thus, we numerically simulate the system within and outside of linearity to confirm Eq. 15 and determine whether these formations survive.

Numerical simulations of M protein density fraction evolution were performed with a finite difference method along a uniform grid (details provided in Finite Difference Method). Figure 4a shows the final frame within linearity for three different simulations at different effective interaction energies, with Fig. 4b displaying a later time image prior to Ostwald ripening dominating the system for the same simulations. While density variations are smoothed out in Fig. 4a, extrema appear roughly periodically, increasing in frequency as *ϵ*_*m*_ increases, agreeing with Fig. 3b. Furthermore, the time needed to reach linearity decreases with increasing effective interaction energy. These trends are confirmed by calculating *q*_*max*_ and *ω*_0,*max*_ for each simulation, as described in Finite Difference Method and shown in S3 Fig. Figure 4c shows these quantities within linearity, where individual simulations for each interaction energy are shown with an empty circle and simulations shown in Fig. 4a are designated with a full circle. With the agreement between simulations and the analytic prediction (colored according to *ϵ*_*m*_, Eq. 24), we confirm that our linear stability analysis holds in linearity.

### High Density Regions Survive Nonlinear Transitions

The numerically determined evolution of M protein density from linearity to nonlinearity involves a sharp transition to a regime of diffusion limited growth where clusters gradually deplete the unclaimed surrounding protein. After depleting the reservoir of isolated protein, these clusters very slowly remodel, where they grow and shrink similarly to Ostwald ripening, typical of Model B dynamics [28]. Furthermore, through comparison between Fig. 4a and 4b, it is clear that maxima within linearity survive cluster formation. This behavior can be seen for a variety of effective interaction energies in S1 Vid-S5 Vid, and for the shown systems in S10 Fig. Note that the small subset of clusters that form during the initial transition and quickly dissipate coincide to weaker maxima within the linear regime. With AFM formations seemingly within the diffusion limited growth regime (Fig. 2d), analysis is only performed on simulations prior to Ostwald ripening dominating the system through a cut-off time. The cut-off time chosen for each simulation is shown above its x-y density plot, as seen in Fig. 4b, with more information in Finite Difference Method and exact cut-off times shown in S5 Fig. Similar to linear stability analysis and Fig. 4a, larger effective interaction energies decrease the time it takes for the system to reach the end of the diffusion limited growth dominated regime.

To quantify the survival of high density regions beyond linearity, the nearest neighbor distance between cluster formations within diffusion limited growth is used in combination with the maximum wavevector determined within linearity. Average nearest neighbor distance (*d*) is shown for each set of effective interaction energies in Fig. 4d, where orange points correspond to simulations shown in Fig. 4a and 4b and black to extra replicates. The analytic prediction, Eq. 16, from linear stability analysis is shown in blue, where the corresponding equation is highlighted. Spread between replicates for a given effective interaction energy are a result of the non-infinite size of the simulated box and the discrete number of clusters. Additionally, the slight overestimate of *d*, particularly for *ϵ*_*m*_ = 9*/*9.25 *k*_*B*_*T*, stems from the initial stage of Ostwald ripening. Refer to S4 Fig for the process of calculating average nearest neighbor distance (*d*) with the images in Fig. 4b. The evolution of this quantity, in addition to the number of clusters and density variance, for each effective interaction energy is shown in S5 Fig, where more information on nearest neighbor distance can be found in Finite Difference Method.

With the agreement in average cluster nearest neighbor distance for linear stability analysis and numerical simulations beyond linearity at a variety of effective interaction energies, Eq. 15 holds beyond linearity. Thus, it can be used to estimate effective interaction energy from AFM cluster formations. In Fig. 4c and 4d, every *ϵ*_*m*_ should have five replicates, where some points are so similar to the point of obscuring each other.

### Effective Interaction Energy from AFM Images

Protein clusters are identified following the filtering and thresholding of the raw AFM data displayed in Fig. 2d and described in Atomic Force Microscopy Cluster Determination. Figure 5a shows the image with the highest area coverage, since clusters are not isotropically distributed otherwise, where red dots represent the centroid of a cluster. A representation of this cluster density for every area coverage is shown in S8 Fig. Binned nearest neighbor distances for each cluster is shown in Fig. 5b, where the red curve represents a kernel density estimate of the histogram and the dotted red line signifies the corresponding maxima, equated to *d* = 77.2 nm. A comparable average nearest neighbor distance for a simulation with *ϵ*_*m*_ = 9 *k*_*B*_*T* is shown through the black dotted line as *d* = 79.6 nm. This simulation is shown on the leftmost panel in Fig. 5c and 5d.

**Fig 5.**
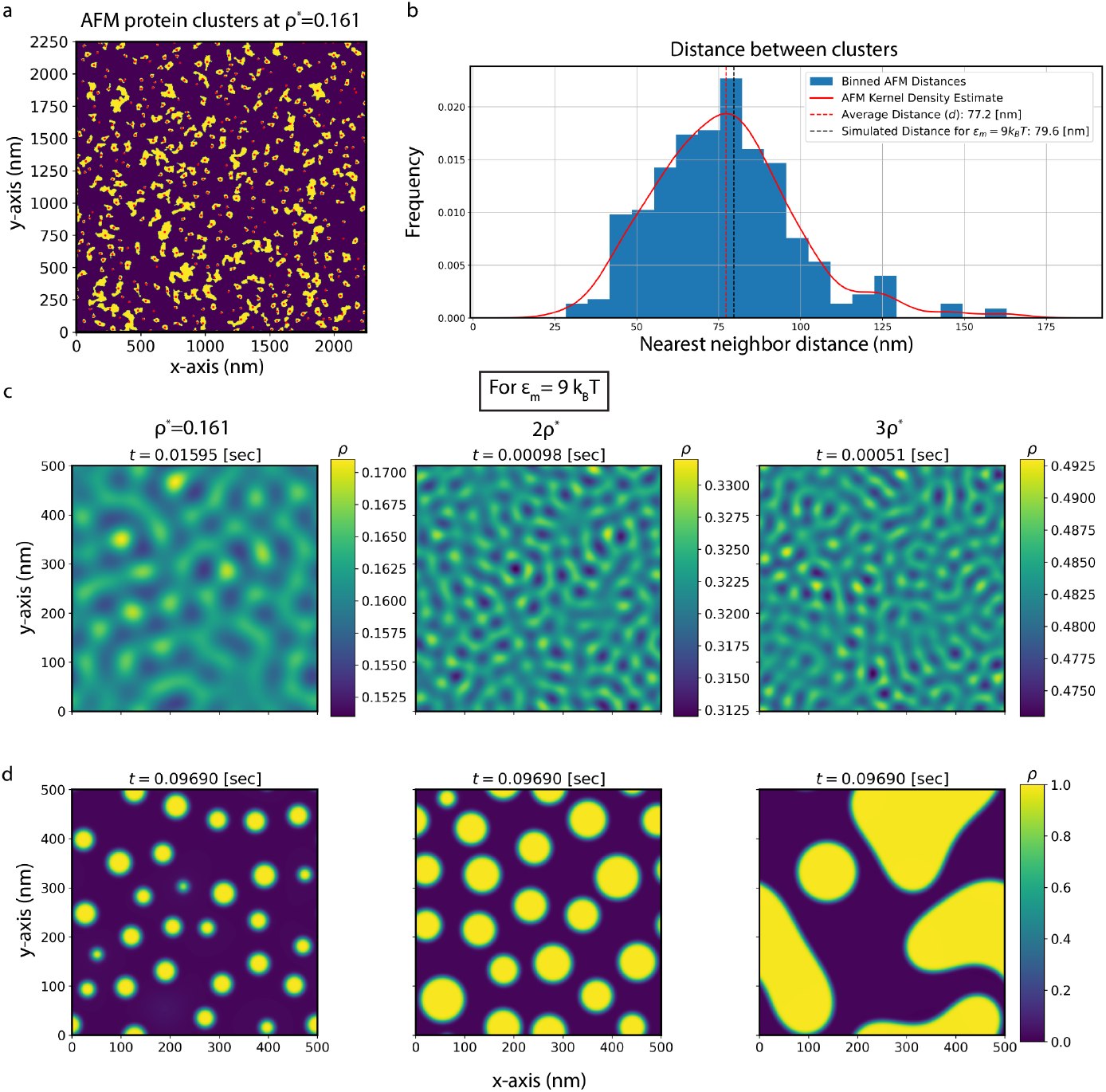
Characterisation of AFM cluster formation and comparison to numerical solutions of the continuum model at a variety of protein density fractions. (a) Filtered and thresholded AFM image at the highest protein area coverage, *ρ*^∗^ = 0.161 ± 0.005. Protein clusters larger than four pixels are shown in yellow with their corresponding centroid in red, while other regions are in purple. (b) A normalized histogram of nearest neighbor distance between protein clusters is shown in blue, where the kernel density estimate is shown in red. The average nearest neighbor distance is taken as the point corresponding to the maximum in the kernel density estimate, and is shown as a red dotted line. A dotted black line displays a comparable *d* for a simulation with *ϵ*_*m*_ = 9 *k*_*B*_*T* . Simulation frames are shown at (c) the final frame of linearity and (d) *t* = 0.0969, with time at the top of each image. This time corresponds to the stable state before Ostwald ripening takes over for *ρ*^∗^ = 0.161. Initial density fraction increases from left to right. Note that the colorbar in (c) changes for each figure. The simulated value shown in (b) is in leftmost column of (c) and (d).

With this nearest neighbor distance, the effective interaction energy can be calculated using Eq. 15. Accounting for the spread in the nearest neighbor distance 77.2 nm ± 1.2 nm, error in calculating area coverage 0.161 ± 0.005 (refer to S7 Fig), and the size of the protein [3.5, 5.5] nm, gives an effective interaction energy in the range of *ϵ*_*m*_ ∈ [7.8 *k*_*B*_*T*, 9.6 *k*_*B*_*T* ].

Next, Fig. 5c and 5d indicate the impact of higher initial protein density in cluster formation within our model. While linearity is still in agreement with our analytic predictions for *ρ*^∗^ = 0.161, as seen in Fig. 5c and S11 Fig, this is not the case for higher densities beyond linearity. For an initial density fraction of 2*ρ*^∗^, shortly after transitioning beyond linearity, clusters begin to merge together. This leads to a much larger distance between clusters than the predicted quantity. In the case of 3*ρ*^∗^, upon leaving linearity, lines of higher density are formed, with the typical cluster formations only appearing at much later time. In both cases, after sufficient time for stabilization, maxima within the linear regime do not coincide with clusters post transition. The transition beyond linearity for the middle and rightmost panels of Fig. 5d involves coarsening beyond diffusion limited growth. Clusters initially form and then merge together, most likely until there is only a single cluster left. These observations can be seen in S3 Vid, S6 Vid, and S7 Vid.

## Discussion and Conclusion

In this paper, M protein cluster formation vital for assembly and budding of SARS-CoV-2 is examined at a multitude of scales. First, as discussed in [27], the long form M protein was not observed in our AFM experiments. This was also confirmed through the relatively substantial structural change during the long form simulation in [27], and structural determination requiring binding with a FAB complex [19, 26]. Whether the lack of long form stems from the absence of additional viral structural proteins or their sole existence in the middle of large clusters is unknown. Thus, cluster formation described in this work only involves the short form, and does not involve other features of viral assembly including the induction of membrane curvature and additional viral structural proteins. However, M protein clusters are still responsible for the onset of assembly, and play a prominent role for the remainder of the assembly and budding process.

Utilizing all-atom MD of an individual short form M protein embedded in an ERGIC-like membrane, the protein induced membrane thinning on the scale of 0.5 nm over 12 nm is quantified. The thinning appears purely elastic, without any noticeable correlation between location and lipid composition. As discussed in [27], this elastic deformation is likely a result of mismatch between the transmembrane domain (∼ 4 nm) and the average thickness of the membrane. This profile is similar to the earlier time profile shown in [27], and agrees with observations from [6]. Furthermore, the thinning induced line tension from this deformation can be approximated as discussed in Thinning Induced Line Tension, acting as a membrane mediated attraction between two nearby M proteins on the scale of 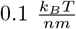 . To determine the role thinning has on M protein cluster formation and to characterize assembly, AFM images of M protein embedded in a suspended membrane are used.

For improved characterization of M protein assembly on the scale of micrometers, these AFM images were gathered at three different protein area coverages, or protein density fractions, within a 2.25 *µm* × 2.25 *µm* membrane. In doing so, the existence of a critical average density fraction in which isotropic clusters form was confirmed, with formations seen at *ρ*^∗^ = 0.161 ± 0.005 but not *ρ*^∗^ = 0.020 ± 0.001. While clusters appeared for *ρ*^∗^ = 0.020 ± 0.001, this can be attributed to large initial local density variations, or aggregation prior to insertion into the vesicle. Smaller oligomers observed within these images could be a result of longer range line tension attracting all nearby protein, eventually reaching a critical distance where no proteins are sensed [63]. Furthermore, clusters of this size/shape and non-periodicity would not prove effective for viral assembly and budding, requiring major remodeling from either M protein spontaneous curvature or interactions with other proteins. These AFM images represent a kinetically trapped state, where M proteins form clusters similarly to diffusion limited growth. This can be related back to [64], where system equilibration was achieved on the scale of minutes for high-speed AFM, much shorter than the scale of our AFM scans. Thus, for a better understanding of M protein cluster formation and viral assembly in a flat plane, these AFM determined final frames are compared to stable states from numerically determining M protein density evolution.

To quantitatively predict M protein cluster formation, we utilized a continuum model describing density evolution in a flat plane. From linear stability analysis, we predicted the existence of a critical density at which clusters form which depends on effective interaction energy. Additionally, for every initial density fraction there exists a critical interaction energy at which clusters form. Using the cluster formation observed in the AFM data, we predicted a lower bound to the effective interaction energy as *ϵ*_*m*_ ∼ 7.4 *k*_*B*_*T* . Furthermore, linear stability analysis provides an estimate of effective interaction energy solely from the average distance between clusters and protein area coverage.

We then used a finite difference method to confirm linear stability analysis for initial density fractions within the range shown in AFM data. In the linear regime, or for little protein density evolution, protein density appears as a smoothed out series of extrema. As these extrema gradually deplete protein from their surroundings they transition to a set of high density clusters in the same location as the initial maxima for low initial density fractions. This behavior is also seen for numerical solutions of a similar model within a low density regime [51]. Within this density regime, linear stability analysis holds in and outside of linearity until later time scale Ostwald ripening dominates the system. For density fractions two or three times higher, linear stability analysis holds within linearity, but does not survive the transition beyond, where clusters coalesce with those around them. This later time evolution is not comparable to the AFM data, which do not display any evidence of coarsening or Ostwald ripening.

Having shown agreement between linear stability analysis and numerical solutions of M protein density, we compared our linear stability predictions with AFM cluster formation to estimate effective interaction energy. From an AFM average nearest neighbor distance between clusters of 77.2 nm ±1.2 nm, we estimated the effective interaction energy to be *ϵ*_*m*_ ∈ [7.8 *k*_*B*_*T*, 9.6 *k*_*B*_*T* ]. Through comparison with our MD determined thinning induced line tension of 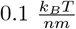, the contribution of line tension to the effective interaction energy is ∼ 1.1 *k*_*B*_*T* (refer to Effective Interaction Energy for more details). Thus, the thinning behavior of short form M proteins does not play a substantial role in cluster formation, and most likely plays a more pronounced role elsewhere in viral assembly and budding. Additionally, direct M-M interactions represent the *ϵ*_*direct*_ ∈ [6.7 *k*_*B*_*T*, 8.5 *k*_*B*_*T* ] left over, with other membrane mediated interactions also possibly playing a role. This relatively strong direct M-M interaction energy is qualitatively in agreement with [16, 17], where a coarse-grained model of SARS-CoV-2 assembly and budding highlights the possible roles of M-M interactions throughout the assembly and budding process [65]. Additionally, the estimated effective interaction energy is comparable to the membrane-mediated protein interactions found in homology predictions of the M protein [66] and for a different transmembrane protein in [64].

Furthermore, confirmation of linear stability analysis outside of linearity for lower density combined with the estimated effective energy provides a density fraction range of [0.118, 0.304] needed for assembly-like cluster formation. In this case, assembly-like clusters are those formed greater than a typical distance of ten protein lengths apart. Even though clusters were capable of forming at lower density in the AFM images, these formations are not indicative of clusters needed for consistent virion assembly. Furthermore, protein interactions within lower density regimes will be dominated by line tension, without the stronger contribution from direct M-M interactions needed for larger cluster formations. For densities substantially larger than this range, it is unclear whether clusters will form consistently as needed for virion assembly, with the possibility of coarsening dominating the assembly process. While this overarching behavior will remain the same, as both curvature and other membrane characteristics (i.e. membrane composition/tension, temperature, suspended membrane, etc.) vary, so to will the provided ranges.

Previous understanding of the assembly and budding process for SARS-CoV-2 did not differentiate the role of the short and long forms of the M protein. Here, the thinning behavior of the short form was confirmed to not play a substantial role in the formation of clusters. Thus, combining this with the work shown from [27], we hypothesize that assembly and budding could start with the formation of short form clusters that transition to long form after successfully recruiting other viral structural proteins along the ERGIC membrane. Upon doing so, the long form coupled to other structural proteins induces curvature in the membrane, possibly improving binding affinity with N/RNA complexes outside the ERGIC, leading to a high protein density membrane bump. With the formation of this dense region of higher curvature, nearby short form will congregate around the edge of the bump, forming a ring capable of thinning the nearby membrane. This ring of thinner membrane surrounding the dense region of higher curvature will then facilitate virion separation from the membrane upon the ERGIC membrane curling around the virus’ genetic material.

Confirming this hypothesis and enhancing the conclusions made in this work requires additional methods at a multitude of scales. The estimated range of interaction energy and the portion of it taken up by direct M-M interactions can be further confirmed utilizing all-atom MD of multiple M proteins embedded within a membrane and performing potential of mean force calculations. Additionally, this method would allow for the determination of conformation dependent protein-protein interactions. To better understand the assembly and budding process, the conformation dependent curvature M proteins induce and those of other structural proteins can be implemented in a continuum model [50–52]. Lastly, a coarse-grained model including these behaviors and combining it with ellipsoidal discrete M proteins would match the non-circular cluster shapes shown in AFM images stemming from either preferential axes of interaction or the shape of the proteins.

Our work suggests that attempts at inhibiting cluster formation and ultimately viral assembly can focus on reducing the binding energy of M-M through mutations (without needing to account for the energy contribution from line tension) such that it falls below the critical effective interaction energy for a given density. Additionally, assembly inhibition can also be achieved through decreasing the amount of M produced such that it falls below the provided range. Ultimately, the improved understanding of M protein assembly and the role of membrane thinning provides insights into alternative methods for preventing viral replication and implications for other enveloped viruses.

## Supporting information

**S1 Fig.**
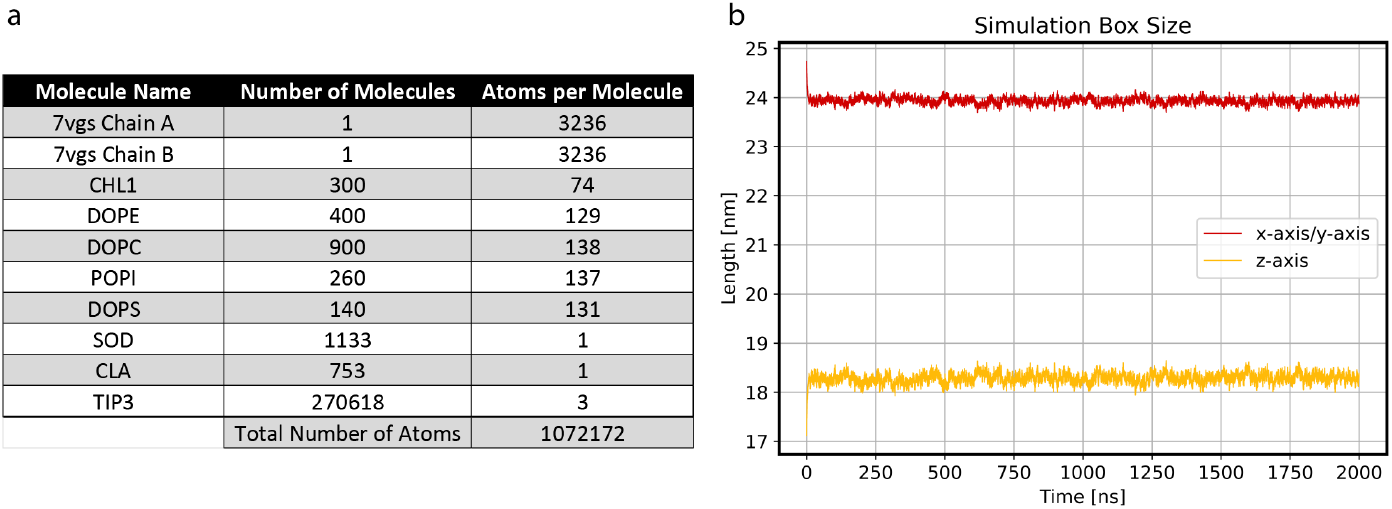
All-atom molecular dynamics atom counts (a) and simulation box size (b). (a) Since the leaflets are symmetric, the number of molecules for a corresponding lipid type in a leaflet is half the system-wide value, adding to 1000 lipid molecules per leaflet. (b) After a quick change in the length along each axis within the first few nanoseconds, the box remains stable throughout the simulation.

## S1 Appendix Nonlinear form of continuum model and conversion to linearity

With our complete free energy shown in Eq. 17, the corresponding variational derivative can be seen in Eq. 18.

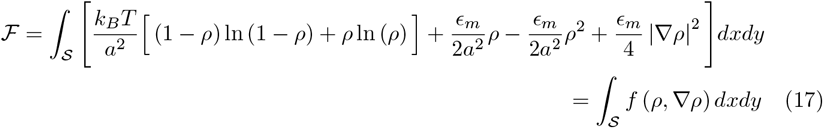

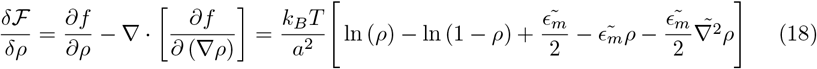

Tildes designate nondimensionalization, as seen with 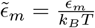 and 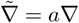 . As a result, the nondimensionalized conservation equation becomes:

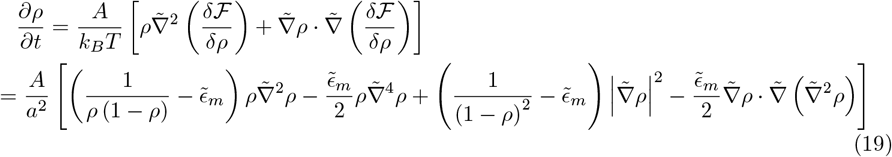

With the form of Eq. 7, where *δρ <<* 1, the following is true and converts Eq. 19 to linearity.

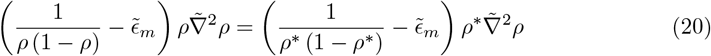

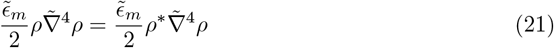

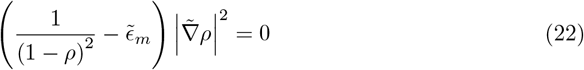

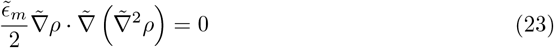

As a result, the linearized density evolution equation can be seen in Eq. 9. Plugging Eq. 8 into the final linear form leads to the dispersion relation shown in Eq. 11. From here, the maximum wavevector (*q*_*max*_) and growth rate (*ω*_0,*max*_) can be defined accordingly for 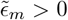.

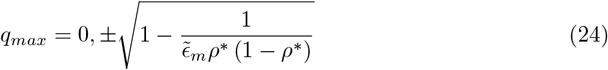

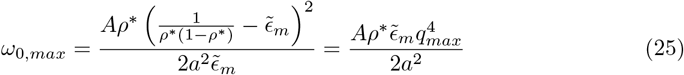

Solving Eq. 24 for 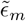 leads to Eq. 15, while using 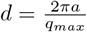 with Eq. 24 leads to Eq. Additionally the curve shown in Fig. 4c and S11 Fig, is Eq. 25 for the corresponding density fraction.

## S2 Appendix Approximating the contribution of line tension in effective interaction energy from a discrete protein lattice to a continuum

To account for the contribution of line tension in effective interaction energy, the energy difference for proteins far apart and close together is determined and compared to the form shown in Eq. 12. As seen in S2 Fig panel a, we start with a square protein lattice, where sites can be occupied with protein or membrane and are a distance *a* apart (where *a* is equivalent to the size of an M protein). Initially, two M proteins are spread apart such that they do not experience any nearest neighbor interactions. The corresponding energy for a general system with only two proteins can be seen below, where *N* is the total number of site-site interactions.

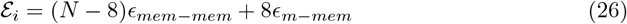

Next, in S2 Fig panel b, these proteins are moved together such that they experience nearest neighbor interactions. As a result, the total energy of the system (ℰ _*f*_ ) is seen in Eq. 27.

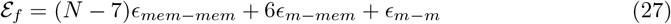

Thus, the energy difference (Δ ℰ_*f*_ ) can be found in Eq. 28, where the factor of *z* is a result of adding the contribution from each axis (z defines the number of nearest neighbors, equal to four with a square lattice).

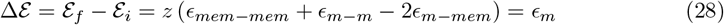

Now, since the energy difference from this discrete lattice is equivalent to the effective interaction energy *ϵ*_*m*_, the energetic benefit of bringing these two proteins together can be equated to this effective interaction energy. Initially, the contribution from line tension for two separate M proteins is shown below, proportional to the surface area, where *γ*_*m*_ is the line tension shown in Eq. 2. Proteins are considered square due to the square lattice chosen in our derivation and numerical solution. The negative sign keeps convention consistent with *ϵ*_*m*_ *>* 0 representing attraction.

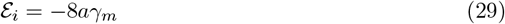

As the square proteins are brought together, the surface area changes to a larger rectangle with a surface area of 6*a*, and direct protein-protein interactions play a more prominent role.

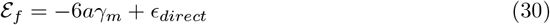

With Eq. 29 and Eq. 30 the energy difference (Δ ℰ) can be determined and compared to the effective interaction energy.

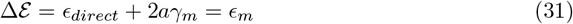

From this method, the contribution from thinning induced line tension in effective interaction energy can be estimated.

**S2 Fig.**
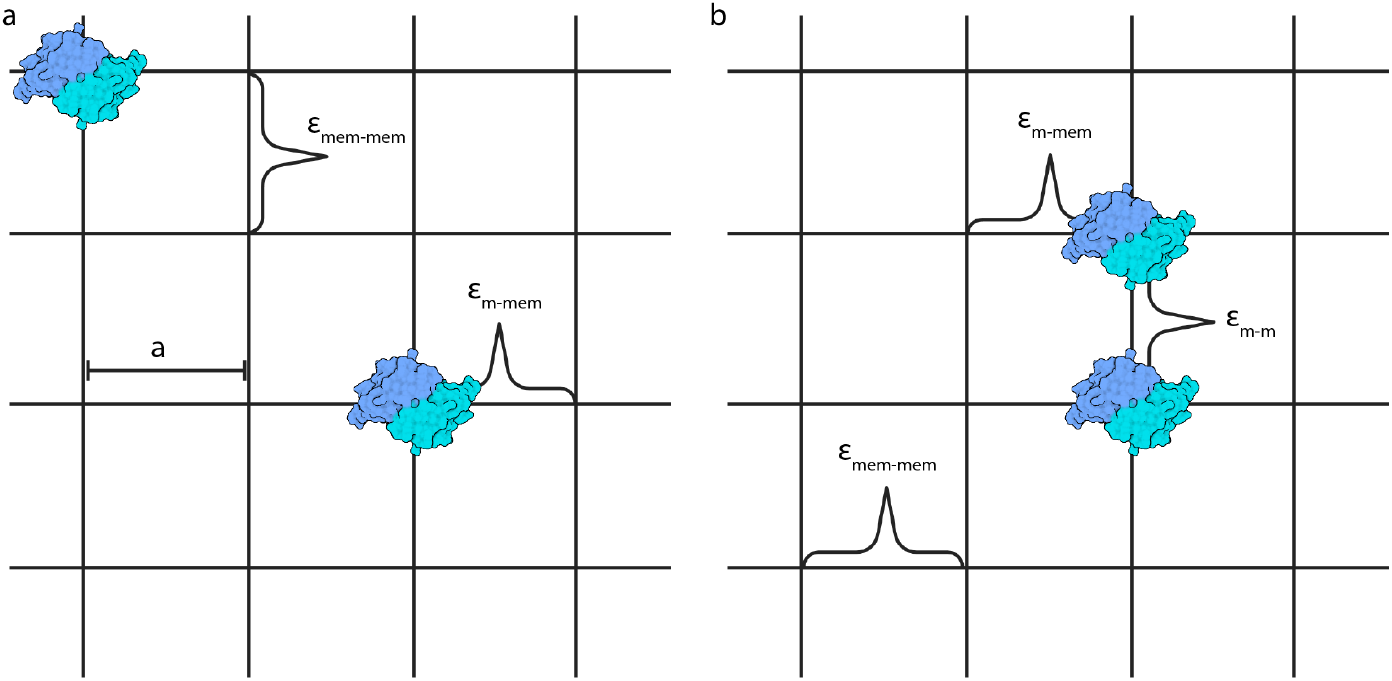
Schematic of a discrete protein lattice with M proteins (a) far apart and within range of (b) nearest neighbor interactions. Discrete protein lattice, with distance between lattice sites equivalent to the size of the protein *a*, shows the two prominent types of site-site interactions when two proteins are not nearest neighbors: *ϵ*_*mem*−*mem*_ and *ϵ*_*m*−*mem*_. (b) As these proteins become nearest neighbors, *ϵ*_*m*−*m*_ encompasses direct interactions between proteins. Converting this representation for a system of randomly distributed proteins to a continuum results in the described continuum model. Created with BioRender.com.

**S3 Fig.**
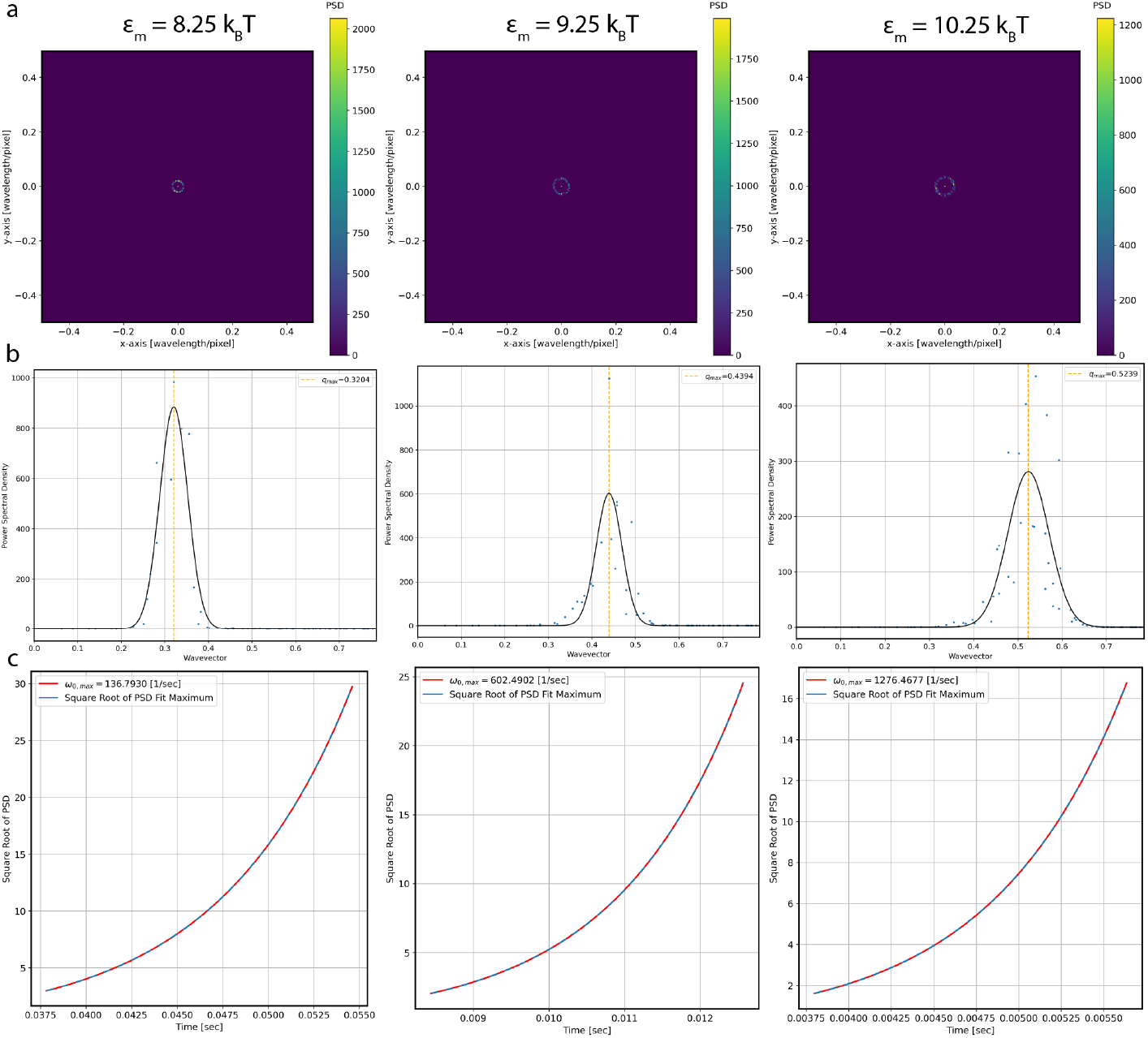
Process for calculating maximum wavevector and growth rate from numerical simulations. (a) Power spectra are shown for the three images displayed in Fig. 4a, where the colorbar displays power spectrum density (PSD) dependent on wavelength per pixel. The radius of the prominent ring is the maximum wavevector. (b) After radially binning from the center of the spectra and averaging the PSD of non-unique radii, the radial profile as a function of the wavevector is shown for each effective interaction energy (wavevector is 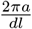 times wavelength per pixel). Blue dots represent the average PSD for every distance from the center, while the black curve shows the corresponding gaussian fit with the maximum wavevector shown as the dotted orange line. (c) The square root of the PSD fit maxima for each measured time is shown with the dotted blue line, while the exponential growth fit is shown in red. The growth rate used for each of these plots is the corresponding maximum growth rate.

**S4 Fig.**
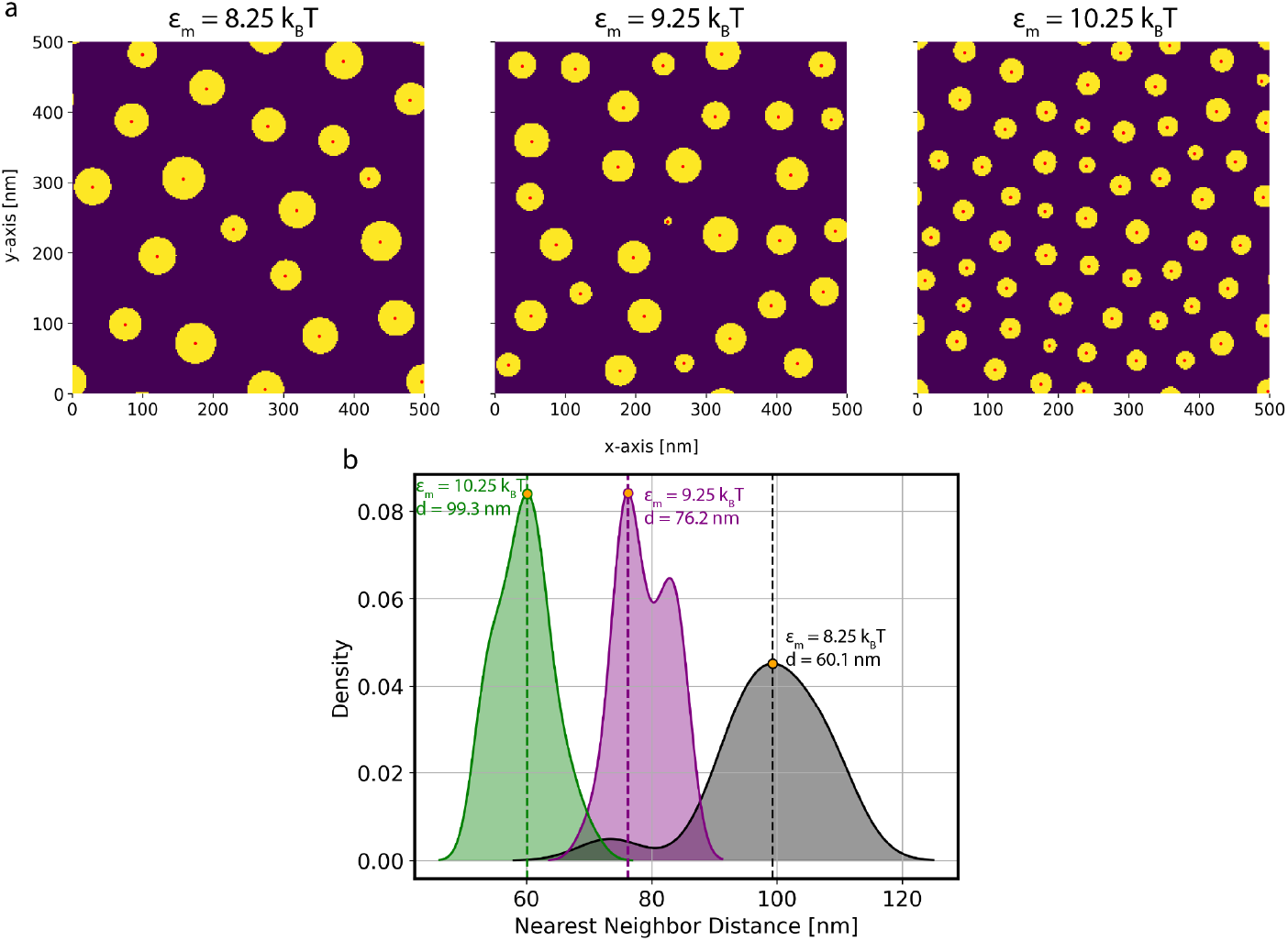
Thresholding (a) and finding nearest neighbor distance (b) for simulated cluster formation. (a) Plot of the thresholded simulations at the cut-off time shown in Fig. 4b, where regions with protein are shown in yellow and cluster centroids are shown in red. (b) Kernel density estimate for each of the three images in black (*ϵ*_*m*_ = 8.25 *k*_*B*_*T* ), purple (*ϵ*_*m*_ = 9.25 *k*_*B*_*T* ), and green (*ϵ*_*m*_ = 10.25*k*_*B*_*T* ), where the dotted line displays the nearest neighbor distance in which there is a maximum (*d*). Each of these distances are shown as orange dots in Fig. 4d.

**S5 Fig.**
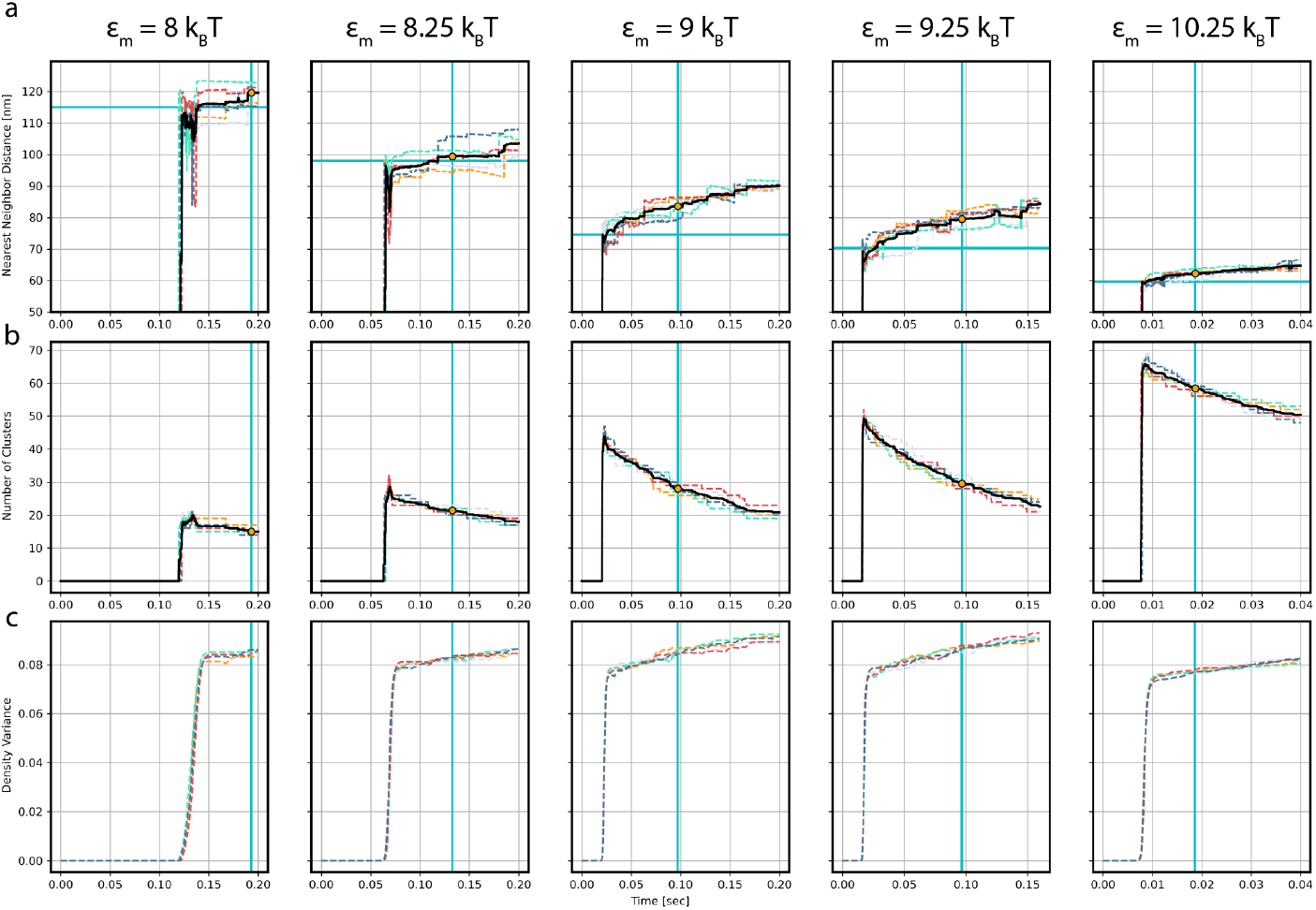
Evolution of simulated nearest neighbor distance (a), number of clusters (b), and variance (c) for every simulation performed at *ρ*^∗^ = 0.161. Each replicate is shown as a dotted line of variable color, where the corresponding average of all replicates is shown as a dotted black line for (a) and (b). Vertical cyan lines represent the cut-off time calculated by finding the time of minimal slope in the average nearest neighbor evolution plot. This time changes for each interaction energy, where *t*_8_ = 0.193 [sec], *t*_8.25_ = 0.13265 [sec], *t*_9_ = .0969 [sec], *t*_9.25_ = .09668 [sec], and *t*_10.25_ = .01863 [sec]. The average value in (a) and (b) at the cut-off time is displayed with an orange dot. For (a), the horizontal cyan line represents the analytically predicted nearest neighbor distance. In (c), the slight fluctuations or bumps in density variance are a result of the gradually decreasing number of clusters stemming from Ostwald ripening shown in (b) for each interaction energy. None of these quantities were gathered until *ρ*_*max*_ *>* 0.9.

**S6 Fig.**
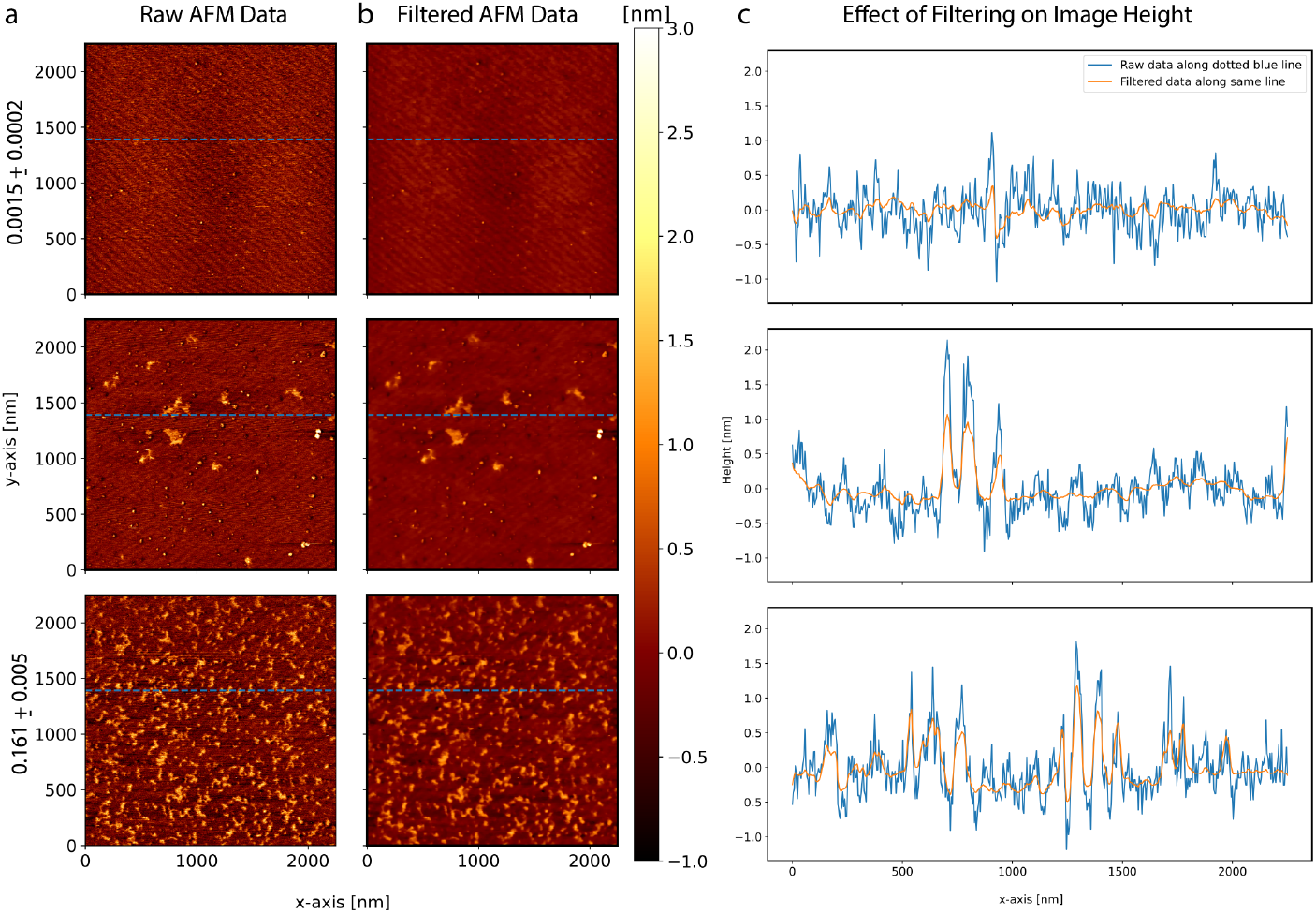
Using a total variation filter to reduce noise of raw AFM data while maintaining large protein clusters. (a) The raw AFM data at the three different protein area coverages shown in Fig. 2a. (b)-(c) This AFM data was filtered with total variation denoising using split-Bregman optimization, with a denoising weght of one and a tolerance of 1 × 10^−5^. Height values for the cross-section shown in (a) and (b) as a dotted blue line is displayed in (c) for comparison between the raw and filtered data.

**S7 Fig.**
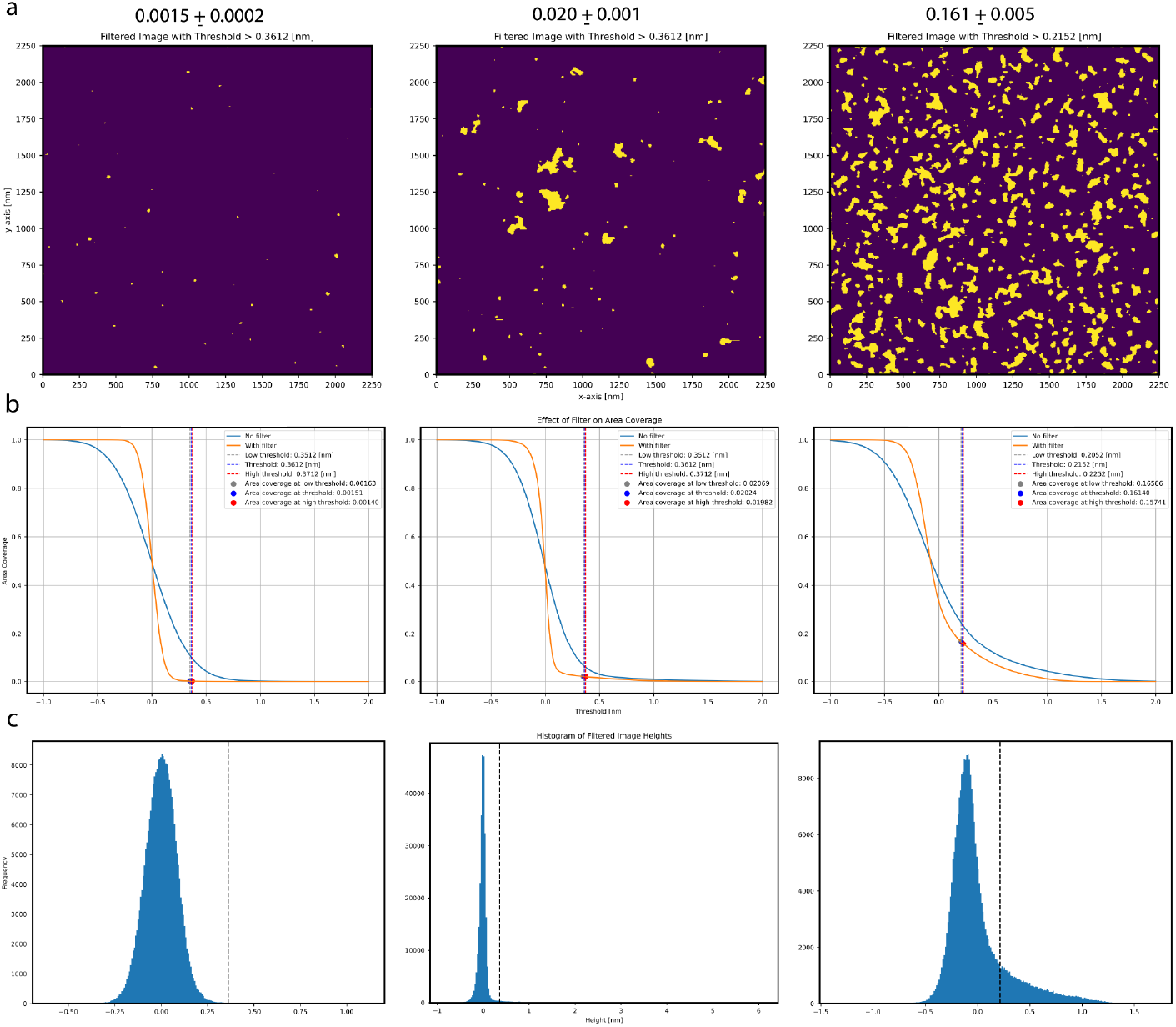
Thresholded and filtered AFM images for the three different protein area coverages. Height thresholds for the two highest area coverages were found using Otsu’s method, where the threshold can be found above each image in (a). Due to the low protein density in the leftmost image, its threshold was chosen to match that of the middle panel.(b) Protein area coverage is shown as a function of threshold height, where the used value is shown as a dotted vertical blue line. As the filter is applied, the curve gets closer to a step function, as expected for a transmembrane protein. The dotted grey line represents the higher bound on area coverage when considering AFM vertical resolution, while the dotted red line signifies the reverse, leading to the error bars in (a). Lastly, (c) displays a histogram of filtered image heights, with the threshold as a dotted black line. Note that Otsu’s method loses effectiveness as the data becomes more Gaussian.

**S8 Fig.**
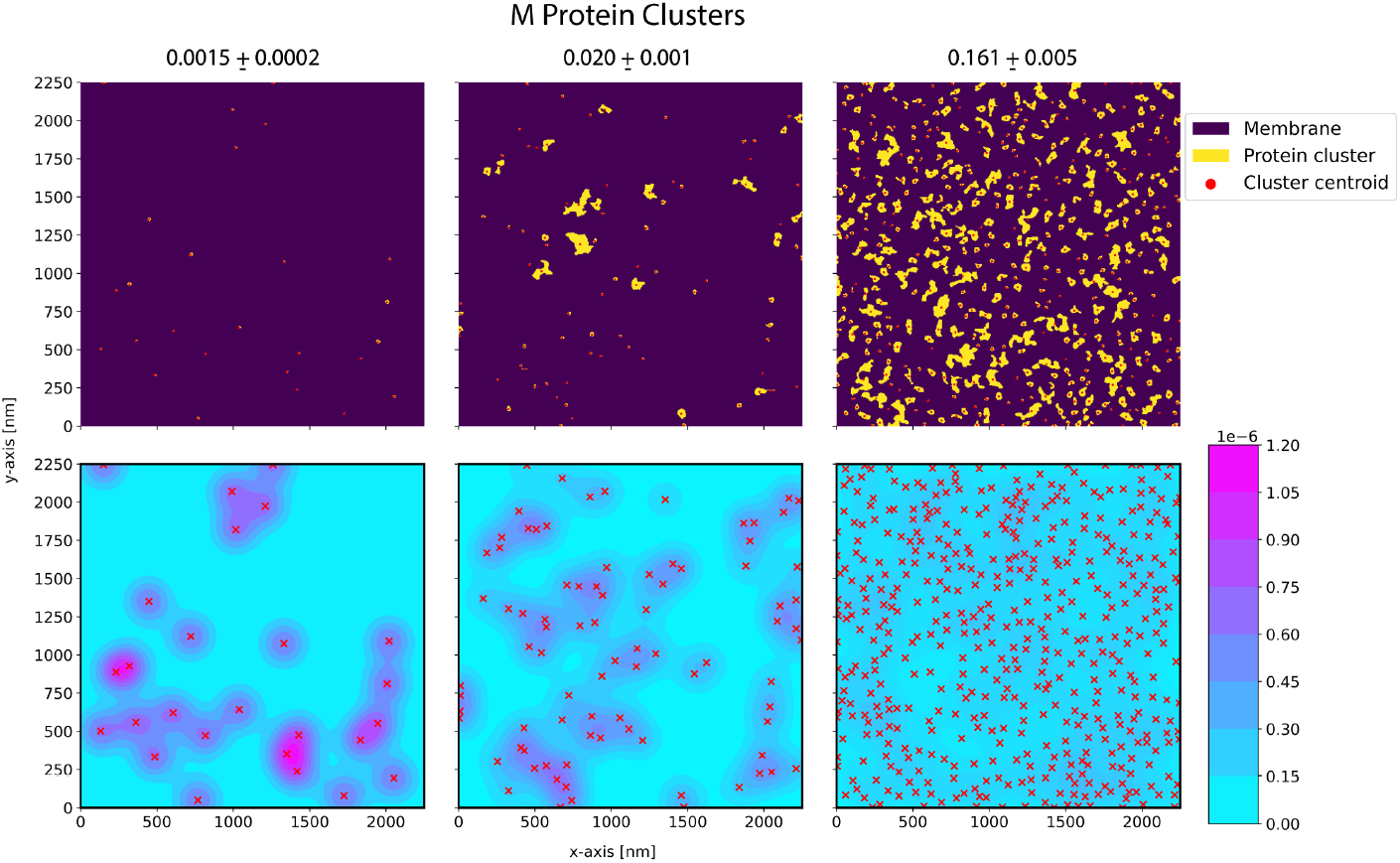
Cluster centroid density for each protein area coverage. The top panel represents filtered and thresholded AFM data for each area coverage with cluster centroids shown as red dots. A cluster is defined as five or more adjacent pixels with height values greater than the threshold. Two-dimensional kernel density estimates of centroids are shown in the panel below, where higher density is shown in pink and lower is shown in blue. Of the three area coverages, only the highest has a close to constant density, showing isotropic cluster distribution.

**S9 Fig.**
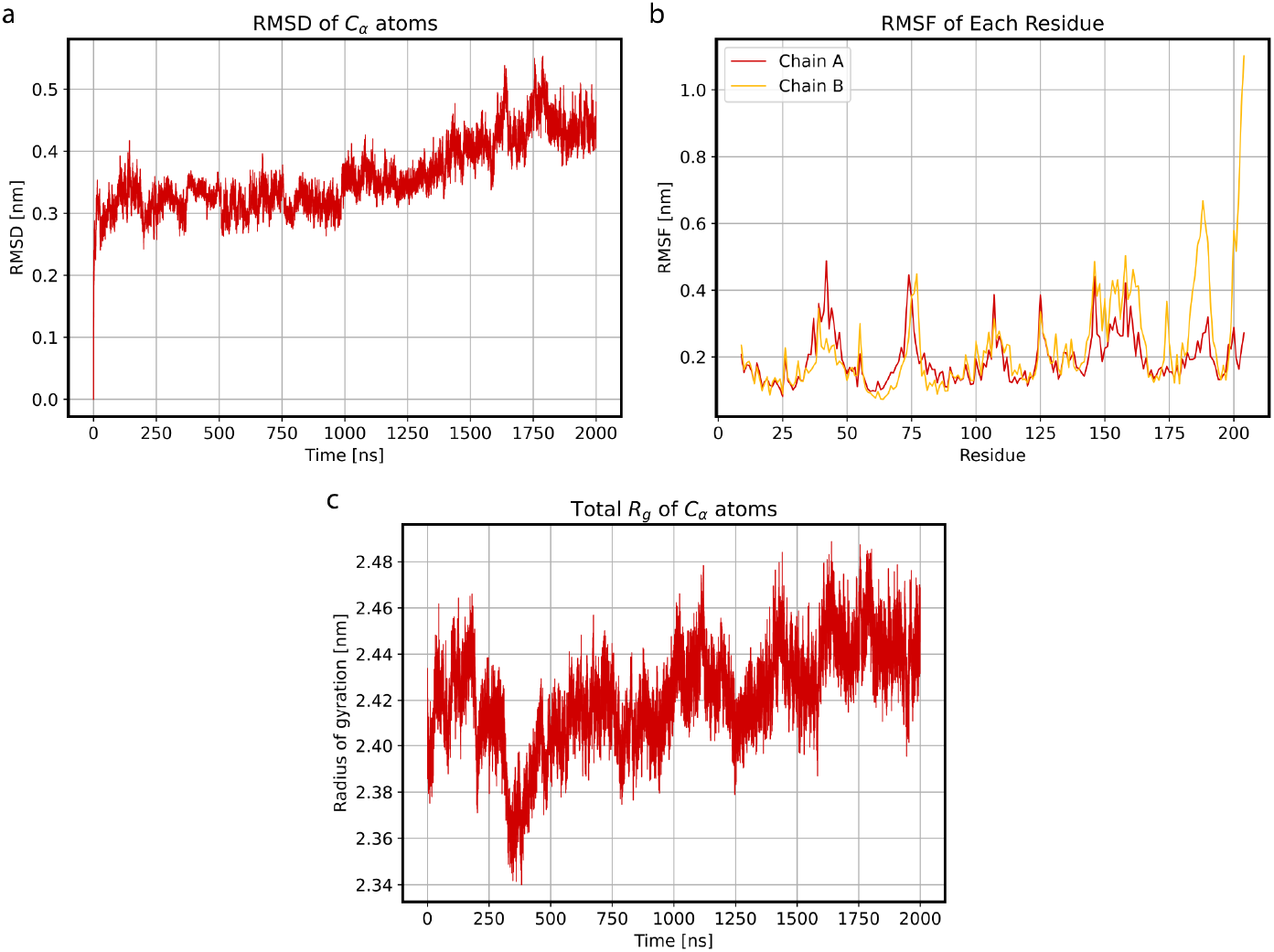
The short form remains stable while embedded in an ERGIC-like membrane throughout the 2 *µs* all-atom MD simulation. (a) Root mean square deviation (RMSD) defines how much a protein structure has changed relative to its initial position. It is defined as 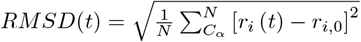, where N is the total number of *C*_*α*_ atoms, *r*_*i*_ (*t*) is the position for the i’th *C*_*α*_ atom at time *t*, and *r*_*i*,0_ is the initial position of the corresponding atom. (b) Root mean square fluctuation (RMSF) is shown for both short form chains, where the N-terminal starts at residue 9 and the C-terminal ends at residue 204. RMSF is the averaged RMSD over time for each residue. (c) Total radius of gyration (*R*_*g*_) of short form *C*_*α*_ atoms is shown in red. With little change throughout the simulation in (a) and (c), and reasonable RMSF for each residue in (b), the protein remains stable throughout the simulation. All of these quantities were calculated using the appropriate GROMACS command.

**S10 Fig.**
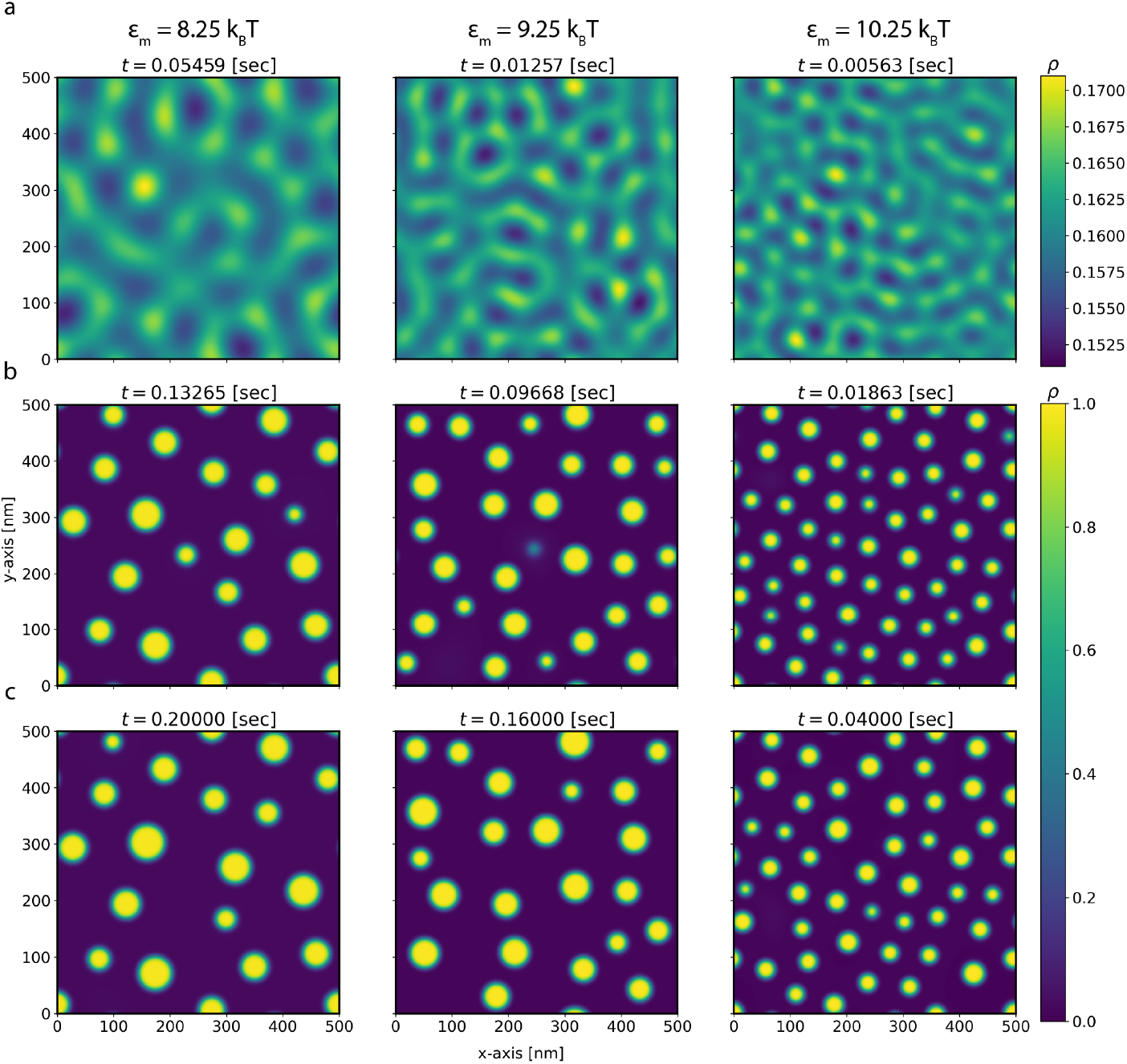
Later time Ostwald ripening shown through density evolution from (a) linearity to (b) end of diffusion limited growth dominated regime to (c) final simulation frame. (a) Frames at the end of linearity from Fig. 4a. (b) Frames before Ostwald ripening dominates from Fig. 4b. (c) Final simulation frames for corresponding interaction energies. Slight dissipation of clusters can be seen in each final frame, where smallest clusters in (b) or weaker maxima in (a) dissipate. Times are shown at the top of each image.

**S11 Fig.**
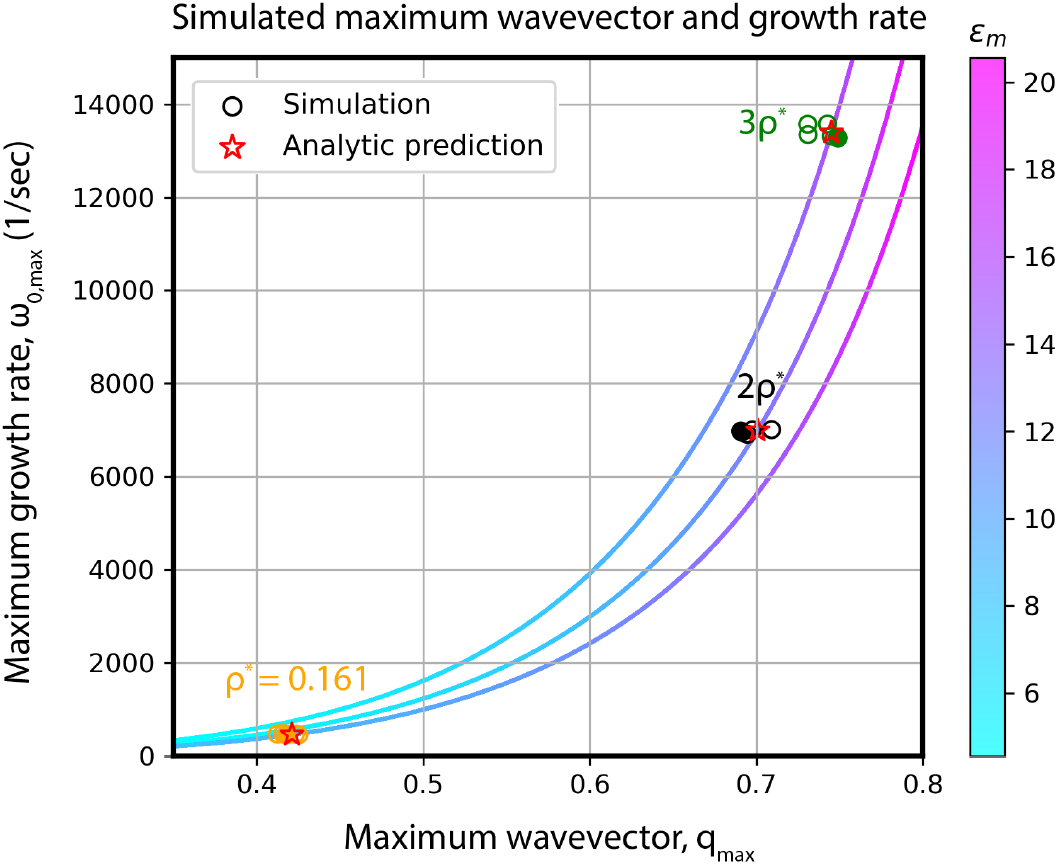
Maximum wavevector and maximum growth rate agrees with predictions within linearity for higher density simulations. Each simulation was performed with *ϵ*_*m*_ = 9 *k*_*B*_*T*, where three five replicate sets at different initial protein densities were used with *ρ*^∗^ = 0.161 shown in orange, 2*ρ*^∗^ shown in black, and 3*ρ*^∗^ shown in green. Filled in circles correspond to replicates shown in Fig. 5. Analytical curves from Eq. 24, where the color gradient represents different effective interaction energies, are shown for each initial density fraction. Red stars are the expected analytic value for the maximum wavevector and maximum growth rate dependent on *ρ*^∗^. The lowest curve corresponds to *ρ*^∗^ = 0.161 at a variety of effective interaction energies, while the middle applies to 2*ρ*^∗^, and the upper 3*ρ*^∗^.

**S1 Vid. Protein density evolution for** *ρ*^∗^ = 0.161 **and** *ϵ*_*m*_ = 8 *k*_*B*_*T* .

**S2 Vid. Protein density evolution for** *ρ*^∗^ = 0.161 **and** *ϵ*_*m*_ = 8.25 *k*_*B*_*T* .

**S3 Vid. Protein density evolution for** *ρ*^∗^ = 0.161 **and** *ϵ*_*m*_ = 9 *k*_*B*_*T* .

**S4 Vid. Protein density evolution for** *ρ*^∗^ = 0.161 **and** *ϵ*_*m*_ = 9.25 *k*_*B*_*T* .

**S5 Vid. Protein density evolution for** *ρ*^∗^ = 0.161 **and** *ϵ*_*m*_ = 10.25 *k*_*B*_*T* .

**S6 Vid. Protein density evolution for** *ρ*^∗^ = 0.322 **and** *ϵ*_*m*_ = 9 *k*_*B*_*T* .

**S7 Vid. Protein density evolution for** *ρ*^∗^ = 0.483 **and** *ϵ*_*m*_ = 9 *k*_*B*_*T* .

